# Melanin Concentrating Hormone- and sleep-dependent synaptic downscaling is impaired in Alzheimer’s Disease

**DOI:** 10.1101/2021.05.21.445007

**Authors:** Sara Calafate, Gökhan Özturan, Nicky Thrupp, Jeroen Vanderlinden, Wei-Ting Chen, Mark Fiers, Ashley Lu, Eline Creemers, Katleen Craessaerts, Joris Vandenbempt, Luuk van Boekholdt, Suresh Poovathingal, Kristofer Davie, Dietmar Rudolf Thal, Keimpe Wierda, Bart De Strooper, Joris de Wit

**Affiliations:** VIB Center for Brain & Disease Research, Herestraat 49, 300 Leuven, Belgium; KU Leuven, Department of Neurosciences, Leuven Brain Institute, Herestraat 49, 3000 Leuven, Belgium; KU Leuven, Department of Otorhinolaryngology, Herestraat 49, 3000 Leuven, Belgium; UK Dementia Research Institute (UK DRI@UCL) at University College London, London WC1E 6BT, UK; Department of Imaging and Pathology, Laboratory of Neuropathology, Leuven Brain Institute, KU-Leuven, O&N IV, Herestraat 49, box 1032, 3000 Leuven, Belgium; Department of Pathology, UZ Leuven, Leuven, Belgium

## Abstract

In Alzheimer’s disease (AD), pathophysiological changes in the hippocampus cause deficits in episodic memory formation, leading to cognitive impairment ^1,2^. Neuronal hyperactivity is observed early in AD ^3,4^. Here, we find that homeostatic mechanisms transiently counteract increased neuronal activity in the hippocampal CA1 region of the *App*^NL-G-F^ humanized knock-in mouse model for AD ^5^, but ultimately fail to maintain neuronal activity at set-point. Spatial transcriptomic analysis in CA1 during the homeostatic response identifies the Melanin-Concentrating Hormone (MCH)-encoding gene. MCH is expressed in sleep-active lateral hypothalamic neurons that project to CA1 and modulate memory ^6^. We show that MCH regulates synaptic plasticity genes and synaptic downscaling in hippocampal neurons. Furthermore, MCH-neuron activity is impaired in *App*^NL-G-F^ mice, disrupting sleep-dependent homeostatic plasticity and stability of neuronal activity in CA1. Finally, we find perturbed MCH-axon morphology in CA1 early in *App*^NL-G-F^ mice and in AD patients. Our work identifies dysregulation of the MCH-system as a key player in aberrant neuronal activity in the early stages of AD.

## Introduction

Despite the early development of neuronal hyperactivity induced by amyloid-β (Aβ) accumulation, Alzheimer’s disease (AD) patients remain cognitively stable for a long time (prodromal phase) ^4,7,8^. This suggests that, before cognitive decline onset, neurons recruit homeostatic plasticity mechanisms to maintain neuronal activity at set-point, within a physiological window ^9, 10^. However, these mechanisms remain uncharacterized. Sleep is essential for memory consolidation ^11^. Synaptic downscaling is a homeostatic plasticity process, prevalent during sleep, that remodels synapses strengthened during wakefulness and removes unnecessary information ^12–14^. Interestingly, AD patients develop sleep disturbances and sleep-silent epileptic-like discharges before any cognitive decline ^15–17^, suggesting impairment of synaptic downscaling. Here we show that initially recruited homeostatic plasticity mechanisms in the CA1 region of *App^NL-G-F^* mice are insufficient to maintain neuronal activity at set-point. Using spatial transcriptomics, we identify Melanin Concentrating Hormone (MCH) as a modulator of homeostatic plasticity. MCH-expressing neurons, which are located in the lateral hypothalamic area (LHA) and project to CA1, are active during sleep ^18,19^ and modulate hippocampal-dependent memory ^6^. We find that the MCH-system becomes dysfunctional in *App^NL-G-F^* mice and in AD patients. Our work provides an important link between dysfunction of the sleep-associated MCH-system, imbalanced homeostatic plasticity, and aberrant neuronal activity in the early stages of AD.

## Results

### Homeostatic plasticity counteracts increased activity of CA1 pyramidal neurons

As expected ^4,20^, we found increased activity in the CA1 region of the hippocampus in *App^NL-G-F^* mice, as evidenced by an increased frequency of spontaneous excitatory synaptic currents (sEPSCs) starting as early as 2 months of age and persisting at 3 months (Fig. 1a), before deposition of Aβ and associated gliosis (Extended Data Fig.1 and 2). While Aβ accumulation was gradual, the increased activity in CA1 strikingly fluctuated over time. Indeed, at 4 months, the frequency of sEPSCs was similar between genotypes, while a transiently decreased sEPSC amplitude (Fig. 1a) and a markedly decreased intrinsic excitability (Extended Data Fig.3) were observed. This suggests that homeostatic mechanisms are recruited to downscale synapses and maintain neuronal activity at a set-point level ^9,21,22^ at this age. However, sEPSC frequency in *App^NL-G-F^* mice strongly increased again at 6 months (Fig. 1a). To independently assess the activity of neurons in the CA1 pyramidal layer over time, we performed immunohistochemistry for the activity-regulated immediate early gene cFos. We observed an increase in both the number of cFos-positive CA1 neurons and cFos signal intensity, followed by a decrease in these parameters at 4 months (Fig.1b and c), mirroring our electrophysiological observations. The cFos signal intensity increased again at 6 months (Fig.1b and c), in parallel with the rising sEPSC frequency (Fig. 1a), suggesting that homeostatic responses are no longer sufficient to maintain neuronal activity at a set-point level at this age (Summarized in Extended Data Fig.7).

**Fig.1.**
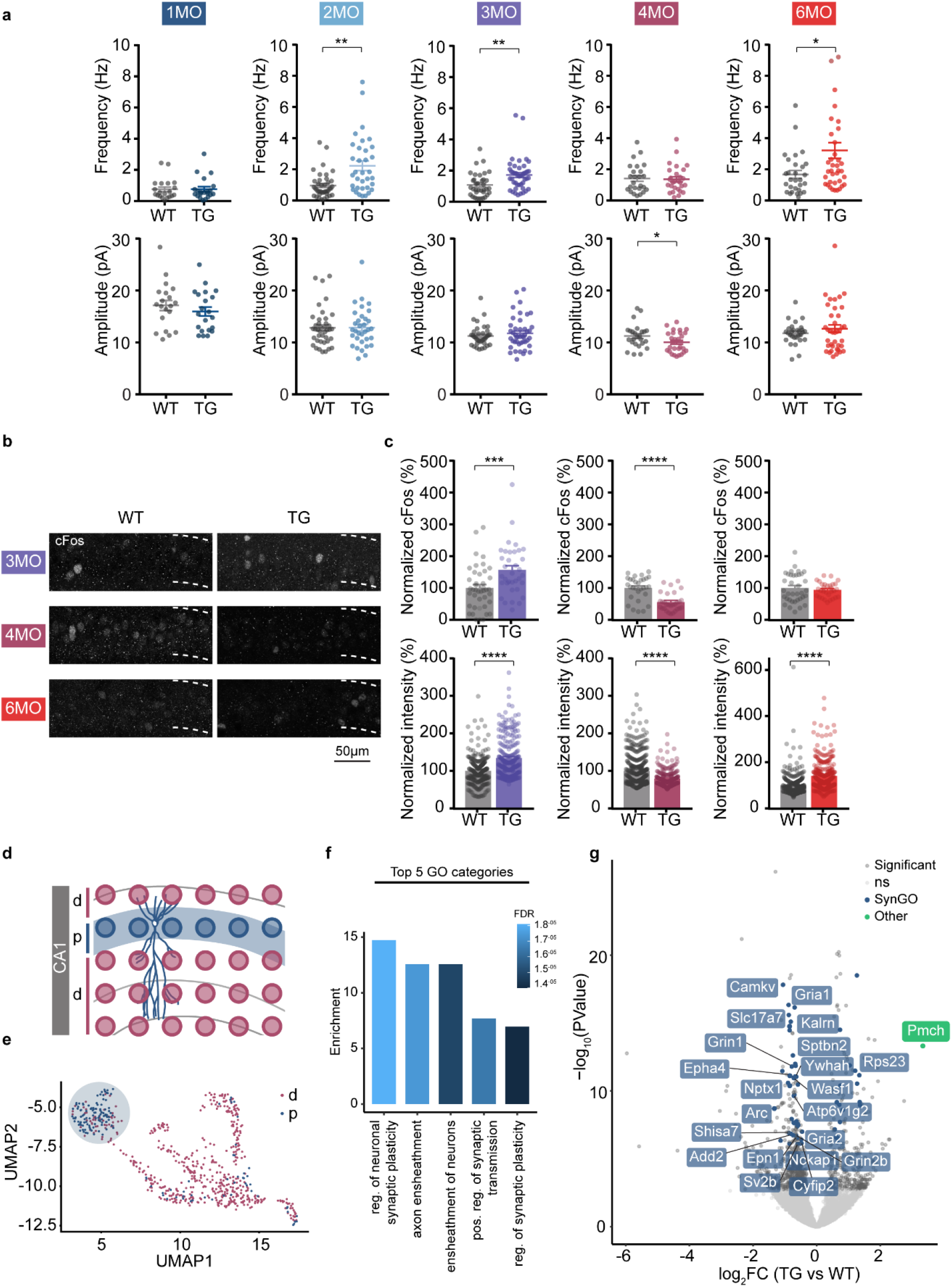
Homeostatic plasticity counteracts increased activity of CA1 pyramidal neurons in *App*^NL-GF^. **a,** Whole-cell voltage clamp recordings of spontaneous excitatory post-synaptic currents (sEPSC) in CA1 pyramidal neurons in acute hippocampal slices from WT and *App*^NLG-F^ (TG) at different months (MO). Number of neurons: n=20, WT and n=21, TG 1MO; n=40, WT and n=34, TG 2MO; n=36, WT and n=44, TG 3MO; n=24, WT and n=28, TG 4MO, n=29, WT and n=38, TG 6MO. Two-tailed unpaired *t*-test: n>3 animals for each genotype. (*P < 0.05, **P < 0.01) **b,c,** Analysis of CA1 pyramidal neuron activity using cFos immunostaining in coronal sections from WT and *App*^NL-G-F^ (TG) mice at 3, 4 and 6 months (MO). **(b)** Representative images of cFos in CA1 neurons at different time points, **(c)** Top graphs show quantification of normalized cFos number of CA1 neurons. Bottom graphs show the normalized intensity of the cFos signal in positive neurons. Number of animals: 3MO - n=6 WT, n=5 TG, unpaired Mann-Whitney test for number and intensity; 4MO - n=5 WT, n=5 TG, Two-tailed unpaired *t*-test for number of neurons and unpaired Mann-Whitney test for intensity; 6MO - n=6 WT; n=5 TG. Two-tailed unpaired *t*-test for number of neurons and unpaired Mann-Whitney test for intensity and twotailed unpaired *t*-test for intensity. (***P < 0.001), ****P < 0.0001). **d,e,** Spatial Transcriptomics (ST) performed on coronal mouse sections of WT and *App*^NL-G-F^ brains at 3.5 months. **(d)** Tissue domains (TD) from CA1 pyramidal layers (p) in blue **(e)** cluster away from dendritic (d) TDs in an unbiased cluster analysis of ST spots from the hippocampus. Number of animals: n=2 WT, n=2 *App*^NL-G-F^. **f,** Results of Gene Ontology (GO) enrichment analysis on the top 200 differentially expressed (DE) ST genes (based on P value). GO categories were sorted by P value and top 8 ordered by normalized enrichment score (Fisher’s Exact Test). 5 most enriched GO categories are shown in the bar plot. Coloring represents False Discovery Rate (FDR). The genes annotated in each GO term are available in Supplementary Table 2. **g**, Volcano plot showing average gene expression differences between *App*^NL-G-F^ (TG) and WT TDs. Significant genes (based on FDR < 0.05, using EdgeR’s quasi-likelihood F test), are shown in dark grey. Significant genes annotated in SynGO are highlighted in blue, other significant genes of interest in green. The 21 genes annotated in SynGO and present in at least one homeostatic plasticity dataset are labelled (Supplementary Table 3). ns=not significant.

### Spatial transcriptomics reveal a homeostatic plasticity signature in the CA1 pyramidal layer

To characterize the homeostatic response of CA1 pyramidal neurons in *App^NL-G-F^* mice at the molecular level, we analyzed our spatial transcriptomics (ST) dataset at 3.5 months of age ^23^. We retrieved small tissue domains (TDs) that cover the pyramidal layer containing cell bodies of CA1 neurons from *App^NL-G-F^* and WT (Fig. 1d and 1e) and analyzed differentially expressed (DE) transcripts between genotypes (Supplemental Table 1). Gene ontology (GO) analysis of the top 200 DE genes (sorted on P value) identified ‘regulation of neuronal synaptic plasticity’ as the top GO category, with 3 out of 5 top enriched GO categories related to synaptic processes (Figure 1f, Supplementary Table 2). Indeed, from the top 200 DE genes, 59 have an annotated synaptic function in the SynGO database (Extended Data Fig.4, Supplementary Table 3). We further cross-referenced the top 200 DE genes with four transcriptomics or proteomics datasets of classical models for homeostatic synaptic plasticity ^24–27^ (Extended Data Fig.5) and found that 62 genes are modulated in at least one of these datasets. From these, 21 genes also have a SynGO annotation (Supplementary Table 3, highlighted in Fig. 1g), including genes such as *Arc*, *Nptx1*, and *EphA4* that are homeostatically modulated upon changes in neuronal activity ^24–26,28^. Thus, consistent with our electrophysiological and immunohistochemical observations, ST analysis reveals a molecular signature of a homeostatic synaptic plasticity response in the *App^NL-G-F^* CA1 pyramidal layer at 3.5 months of age.

### Persistent recruitment of homeostatic plasticity does not compensate aberrant neuronal activity

Prolonged network excitation has been shown to induce persistent homeostatic adaptations in spine density and Hebbian plasticity that can lead to imbalanced synaptic functions ^22,29^. In agreement, we observed a persistent decrease in spine density in proximal apical dendrites from *App^NL-G-F^* CA1 pyramidal neurons that started at 4 months and became pronounced at 6 months (Extended Data Fig.7), as well as an increased threshold for LTP at Schaffer collateral (SC)-CA1 synapses at 6 months (Extended Data Fig.8).

Strikingly, at 6 months of age, a number of CA1 pyramidal neurons appear to bypass activity-dependent homeostatic mechanisms, as illustrated by increased sEPSC frequency and increased cFos signal intensity (Fig. 1a-c). Furthermore, we found an increased AMPA/NMDA ratio of evoked EPSCs at SC-CA1 synapses at 6 months (Extended Data Fig.9), suggesting an increased AMPA receptor (AMPAR) content at these synapses. This suggests that in response to continuous Aβ build-up, mechanisms become imbalanced and fail to maintain neuronal activity at set-point. To determine whether we could detect a molecular signature of an imbalanced homeostatic response, we analyzed the protein levels of several DE genes from the ST analysis in hippocampal synaptosome extracts and tissue sections from *App^NL-G-F^* and WT mice at 6 months (Extended Data Fig.10). We observed an increase in synaptic EphA4 levels in *App^NL-G-F^* synaptosomes, consistent with a persistent decrease in spine density ^28^. Increased Copine6 and decreased Icam5 levels on the other hand correlate with more mature synapses ^30^ (Extended Data Fig.10). Moreover, the increased intensity of Nptx1 immunoreactivity in the CA1 region in *App^NL-G-F^* mice (Extended Data Fig.10) suggests a stabilization of AMPARs ^31^ in remaining synapses. Additionally, decreased synaptic Arc protein levels in *App^NL-G-F^* synaptosomes suggest increased AMPAR function and reduced synaptic downscaling ^32^. To further analyze this, we determined the protein levels of Homer1a and mGluR5, two synaptic molecules that stabilize AMPARs during sleep-dependent synaptic downscaling ^14^. We observed increased mGluR5 and decreased Homer1a levels in *App^NL-G-F^* hippocampal synaptosomes (Extended Data Fig.10), pointing to a stabilization of AMPARs in remaining synapses ^14^. Together, these biochemical and immunohistochemical observations are consistent with an excessive removal of spines, but an increased strength of remaining synapses in *App^NL-G-F^* mice at 6 months, in line with the increased AMPA/NMDA ratio (Extended Data Fig.9). In conclusion, while the homeostatic response to continuous Aβ accumulation is persistent, this is not sufficient to maintain stability of neuronal activity at set-point over time.

### MCH regulates synaptic downscaling

The advantage of the ST approach is that it covers cellular niches in the brain, yielding the transcriptional profiles of CA1 pyramidal neurons but also of the surrounding environment ^23^. ST analysis revealed that the most up-regulated gene in the CA1 pyramidal region of *App^NL-G-F^* is *Pmch*, which belongs to the most enriched GO category (regulation of neuronal synaptic plasticity) (Fig. 1f and 1g, Supplementary Table 2). *Pmch* encodes the prepro-melanin-concentrating hormone peptide (ppMCH) that is further processed into several peptides including MCH ^33,34^. While no precise synaptic function has been attributed to this neuropeptide, MCH-positive hypothalamic neurons projecting to the dorsal hippocampal CA1 region play a role in modulating memory ^6^. Neuronal cell bodies that express *Pmch* mRNA are exclusively located in the LHA ^34^, confirmed by single-molecule fluorescent *in situ* hybridization (Fig. 2a,b and Extended Data Fig.11a,b). CA1 pyramidal neurons from WT and *App^NL-G-F^* do not express *Pmch* (Fig. 2a, Extended Data Fig.11c), but express *Mchr1*, which encodes MCH receptor 1 (MCHR1), the only known receptor for this neuropeptide ^35^. MCH-positive neurons (Fig. 2c) project their MCH-rich axonal terminals from the LHA to the dorsal CA1 region, as shown by MCH immunohistochemistry in sections of adult WT mice (Fig. 2d). This suggests that *Pmch* detected in the CA1 ST analysis is derived from mRNA pools in MCH-positive axons.

**Fig. 2.**
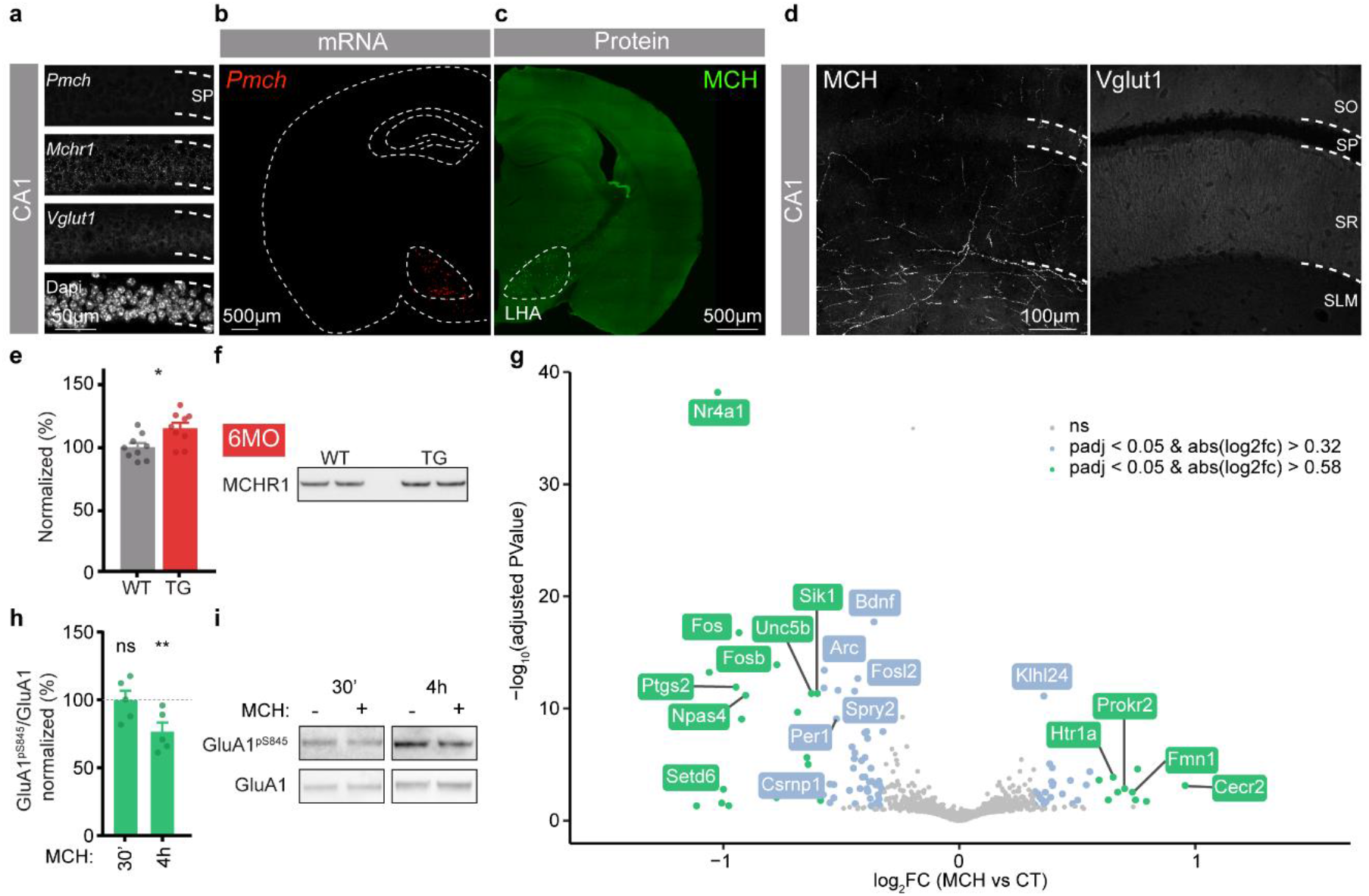
MCH induces a molecular signature of synaptic plasticity in hippocampal neurons. **a,** CA1 pyramidal layer crops showing *Pmch, Mchr1* and *Vglutl* mRNA expression using RNAscope. DAPI was used to visualize nuclei in CA1 pyramidal layer. **b, c,** whole brain coronal section showing that cell bodies expressing **(b)** *Pmch* detected by RNAscope on 10μm section and **(c)** MCH protein detected by immunostaining on 50μm section, are located in the LHA. **d,** CA1 coronal sections showing the presence of MCH-positive axons projecting from LHA. Vglut1 immunostaining was used to identify CA1 laminas. **e, f,** Quantification **(e)** and representative western blot **(f)** for MCHR1 in hippocampal synaptosome preparations from WT and *App*^NL-G-F^ (TG) mice at 6 months. Two-tailed unpaired *t*-test: n=9 animals for each genotype. (*P < 0.05) **g,** Volcano plots showing gene expression differences between average gene expression from bulk RNAseq on hippocampal primary cultures treated with control vehicle (H2O) or 1μM MCH peptide for 4 hours. Number of independent cultures: n=2, 2 replicas per culture. (log2fc values calculated using DESeq2, p-values adjusted using the Benjamini & Hochberg correction). DE genes are available in Supplementary Table 5. **h, i**, Hippocampal primary cultures treated with control vehicle (H2O) or 1μM MCH peptide for 30 minutes or 4 hours were lysed and phosphorylated GluA1 on serine 845 (GluA1^pSer845^) and total GluA1 were detected by western blot. (h) GluA1^pSer845^/GluA1 ratio normalized to control vehicle and (i) representative western blot. Number of independent cultures for all conditions: n=5, data points represent the average of 3 replicas per each independent culture. Two-tailed unpaired *t*-test (**P < 0.01).

How MCH affects CA1 pyramidal neurons to modulate CA1-dependent memory is unknown. We detected MCHR1 in hippocampal synaptosomal extracts and found increased synaptic levels of this receptor in *App^NL-G-F^* mice at 6 months (Fig. 2e,f). This suggests that MCH-MCHR1 signaling at synapses is altered in *App^NL-G-F^* mice. To uncover the neuronal responses to MCH, we treated dissociated hippocampal cultures with MCH peptide for 4 hours and performed bulk RNA sequencing (Fig. 2g and Supplementary Table 5). Strikingly, MCH treatment consistently downregulated several activity-regulated immediate early genes involved in synaptic plasticity: *Nr4a1 (*the most downregulated gene), *Fos, Fosb, Fosl2, Arc* and *Npas4* ^32,36–38^ (Fig. 2g and Supplementary Table 3). Additionally, genes involved in neuronal damage response (*Prokr2*), implicated in epilepsy (*Sik1*), and genes that regulate kainate-mediated responses (*Spry2, Ptgs2* and *Klhl24*) ^39–43^ are also modulated upon MCH treatment (Fig. 2g and Supplementary Table 5). Moreover, genes involved in synaptic transmission and plasticity (*Bdnf, Htr1a*) are also among the top MCH-modulated genes. Together, these observations suggest that MCH downregulates the expression of synaptic plasticity-related genes. To assess the effect of MCH treatment on synaptic downscaling, we analyzed the levels of phosphorylated GluA1 on serine 845 (pS845), a phospho-epitope on the GluA1 AMPAR subunit that promotes GluA1 surface targeting or retention. We found that GluA1 pS845 levels were decreased in total lysates upon 4 hours of MCH treatment (Fig. 2h), suggesting that MCH downscales synapses by reducing surface availability of AMPARs. Altogether, these observations suggest that MCH neurons that project to the hippocampus are implicated in regulating synaptic plasticity and synaptic downscaling of CA1 pyramidal neurons.

### MCH-neuron activity during rebound sleep is impaired in *App^NL-G-F^* mice

Our findings that *Pmch* mRNA levels are altered in CA1 at 3.5 months (Fig. 1g) hint at impaired MCH neuron function. Importantly, Aβ plaques accumulate in the LHA already at 3 months (Extended Data Fig.12). We thus asked whether the activity of MCH neurons was affected in *App^NL-G-F^* mice when compared to WT at 6 months. MCH neurons are active during REM sleep ^44^ and co-distribute with orexin-positive neurons, which promote awakening ^45^. We quantified the activity of these two populations in WT and *App^NL-G-F^* mice at 6 months using cFos expression across 3 different states: at the beginning of the light cycle (awake control group - W), after 6h of sleep deprivation (SD group) and 6h of SD followed by 4h of rebound sleep (SD + RB group) (Fig. 3a). The time spent in NREM and REM sleep increases during periods of rebound sleep and so does the activity of MCH-positive neurons ^19^. We found a similar percentage of active (cFos-positive) orexin neurons between genotypes in all groups, with the majority of active neurons being orexin-positive as the animals are awake (Fig. 3b,c,d). The activity of MCH neurons in the W and SD groups was similar between genotypes (Fig. 3b,d). As expected, SD + RB increased the percentage of active MCH neurons in WT mice. However, the increase in the percentage of cFos-positive MCH neurons in *App^NL-G-F^* mice in the SD + RB group was strongly attenuated compared to WT (Fig. 3b,d). These results suggest that the activity of MCH neurons is impaired in the early stages of Aβ pathology.

**Fig. 3.**
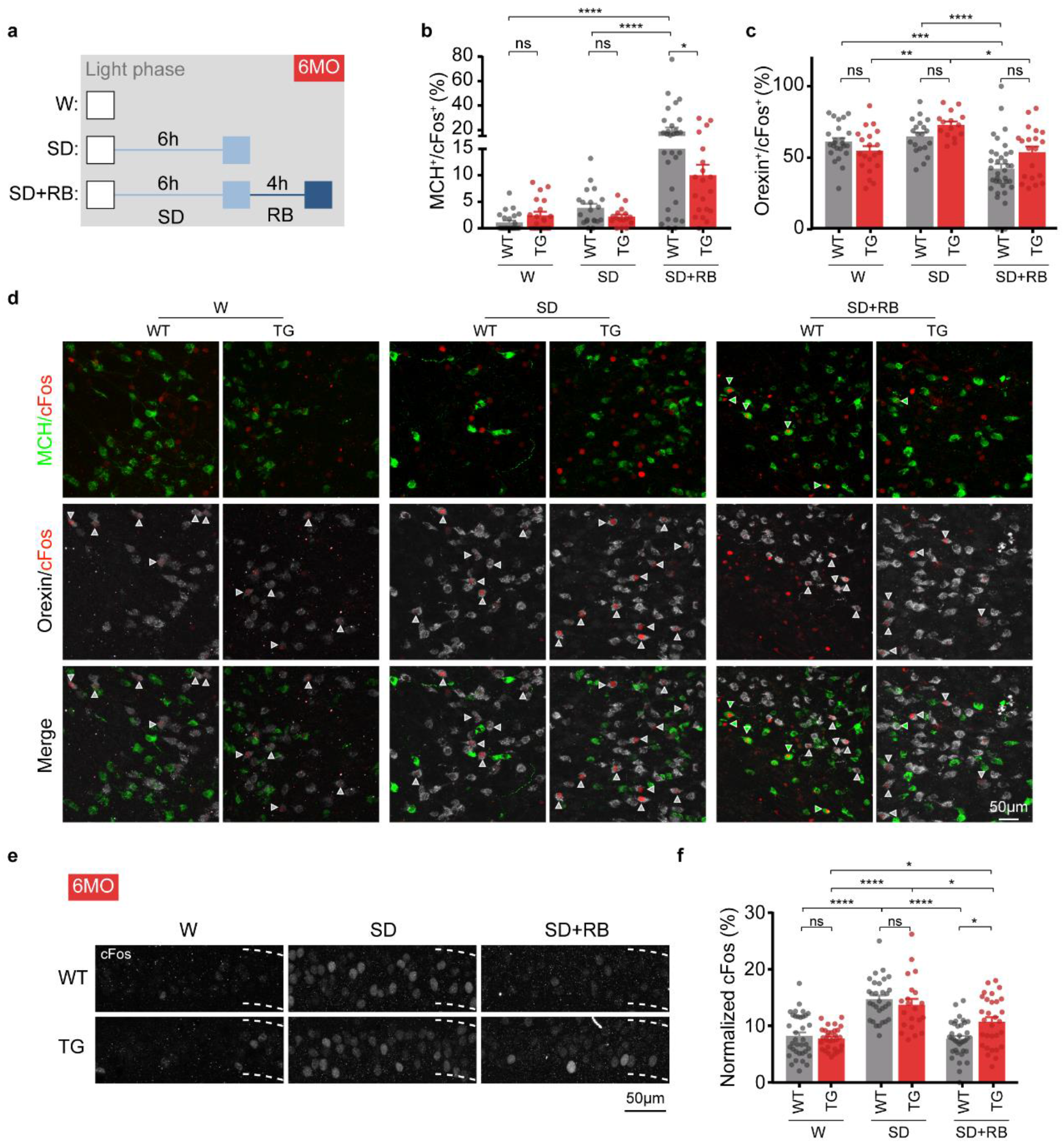
SD reveals impaired activity of LHA and CA1 pyramidal neurons in *App*^NL-G-F^. **a,** Schematic representation of the animal groups used at 6 months. Experiment started at the beginning of the light cycle with 3 paradigms: awake control group (W), 6h of sleep deprivation group (SD) and 4h of rebound sleep following 6h of SD (SD + RB). **b, c,** Quantifications of LHA neurons from WT and *App^NL-G-F^* (TG) mice of different groups (b) MCH^+^/cFos^+^ and (c) Orexin^+^/cFos^+^. W: n=6 WT, n=5 TG; SD: n=5 WT, n=4 TG, SD + RB: n=6 WT, n=5 TG. 3 or more LHA fields analyzed per animal. one-way ANOVA with Tukey’s post-hoc test (*P < 0.05). **d,** Representative images of LHA showing MCH (green) or Orexin (grey) neurons double positive for cFos (red), with green arrows indicating MCH^+^/cFos^+^ and grey arrows Orexin^+^/cFos^+^. **e, f,** Analysis of CA1 pyramidal neuron activity using cFos immunostaining in coronal sections from WT and *App*^NL-G-F^ (TG) mice of different groups. (e) representative image and (f) quantification of the percentage of cFos-positive neurons and intensity of cFos signal. Number of animals W: n=6 WT, n=5 TG; SD: n=5 WT, n=4 TG, SD + RB: n=6 WT, n=5 TG. 3 or more CA1 fields analyzed per animal. one-way ANOVA with Tukey’s post-hoc test (**P < 0.01, ***P < 0.001, ****P < 0.0001).

### CA1 pyramidal neurons in the *App^NL-G-F^* show reduced capacity for downscaling upon sleep deprivation

Neuronal firing rates decrease during sleep ^12^ and SD prevents downward firing rate homeostasis ^13^. Given that the activity of MCH neurons is impaired in *App^NL-G-F^* mice, we hypothesized that a SD-induced increase in CA1 neuronal activity would fail to normalize after rebound sleep in *App^NL-G-F^* mice. As expected, upon SD the percentage of active neurons in the CA1 pyramidal layer increased in both WT and *App^NL-G-F^* mice (Fig. 3e,f). In WT mice, the number of active neurons renormalized after rebound sleep. In contrast, in *App^NL-G-F^* mice the number of active neurons failed to return to basal levels in the SD + RB group (Fig. 3e,f). This result shows that the activity of neurons in the CA1 pyramidal layer fails to return to set-point after rebound sleep, a period when MCH-neurons are active.

### CA1 MCH-positive axons are dystrophic in *App^NL-G-F^* mice and in AD patients

We explored morphological properties of MCH-positive axons in *App^NL-G-F^* and WT mice in the CA1 region. To visualize MCH-positive axons, which contain neurotransmitter release sites ^46^, we used thick sections for immunohistochemistry. We found that the number of MCH-positive puncta per axon length was similar between the two genotypes, but that their area was increased in 6 month *App^NL-G-F^* mice compared to WT mice (Fig. 4a,b). MCH-positive axons in the vicinity of Aβ plaques, revealed by 6E10 immunostaining, adopted a dystrophic appearance, in particular when passing through a plaque ^47^ (Fig. 4c, Extended Data Fig.13). These morphological alterations were detected in the CA1 region already at 3 months of age, even before sleep disturbances appear in these mice ^48^, and were continuously observed as plaques are gradually formed (Extended Data Fig.14).

**Fig. 4.**
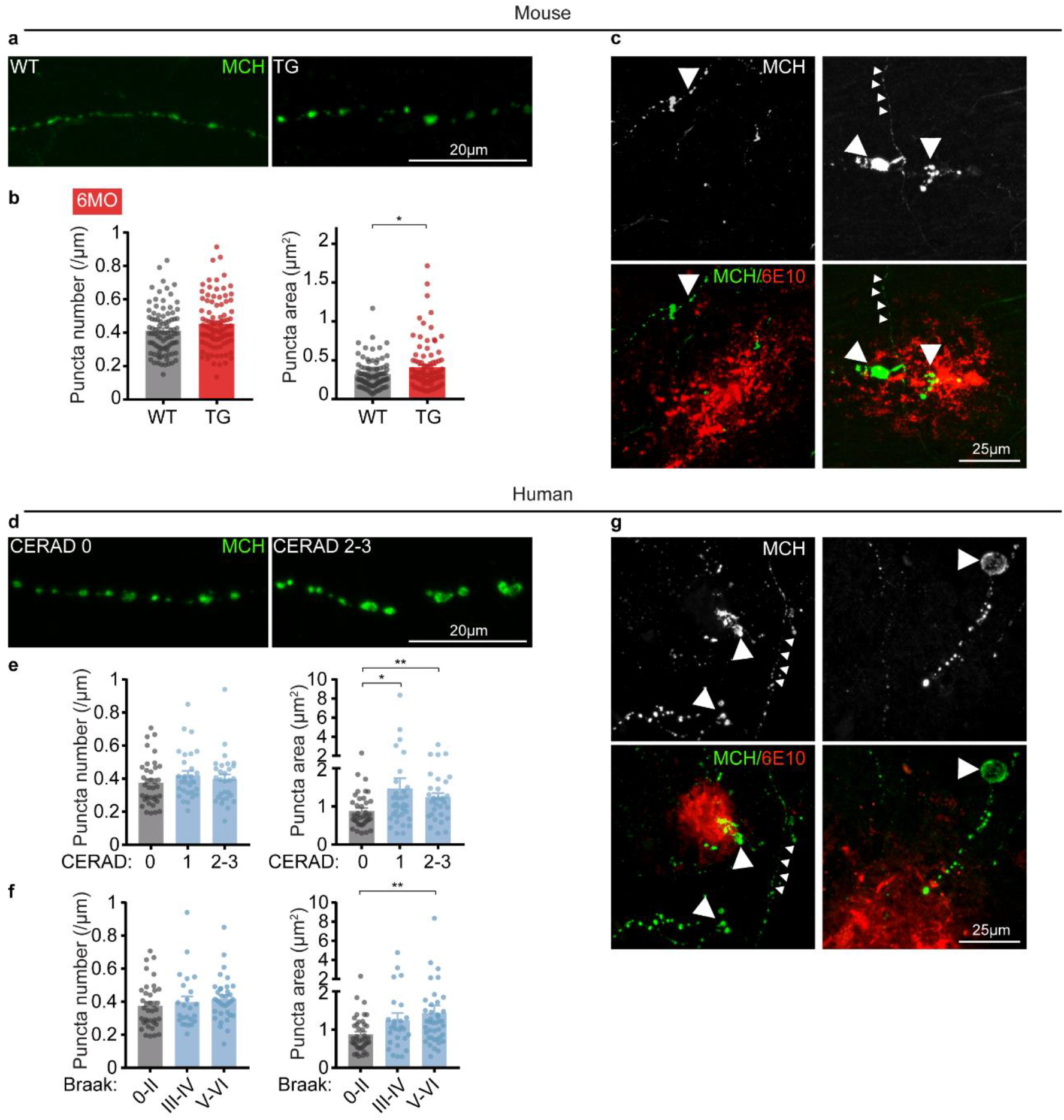
CA1-projecting MCH-positive axons are dystrophic in *App*^NL-G-F^ and AD patients. **a,b,** CA1-projecting MCH-positive axons in WT and *App*^NL-G-F^ (TG) mice at 6 months. **(a)** representative image and **(b)** quantification of the number of MCH puncta along axon length and respective puncta area. Number of animals: n=6 WT, n=5 TG, Two-tailed unpaired *t*-test (*P < 0.01). **c,** Representative images of MCH axons in CA1 hippocampal region showing dystrophic morphology near Aβ plaques labelled by 6E10 antibody (indicated by large arrowheads) and neurite with normal morphology (indicated by small arrowheads), in *App*^NL-G-F^. **d-f,** CA1-projecting MCH positive axons in controls and AD patients at different stages of AD pathology. **(d)** representative image and quantification of the number of MCH puncta along axon length and respective puncta area in relation to **(e)** CERAD and **(f)** Braak stages. Number of patients: n=6 CERAD 0; n=2 CERAD 1; n=3 CERAD2-3; n=6 Braak 0-II; n=2 Braak III-IV; n=3 Braak 0-V.VI; unpaired Mann-Whitney test (*P < 0.05, **P < 0.01). **g,** Representative images of MCH axons in CA1 hippocampal region showing dystrophic morphology indicated by the large arrow near Aβ plaques labelled by 6E10 antibody and neurite with normal morphology indicated by smaller arrows, in one AD patient brain of CERAD score 1.

To determine whether these observations in *App^NL-G-F^* mice are relevant for human AD patients, we prepared thick hippocampal sections from 12 post-mortem human patients at different stages of AD pathology and performed immunohistochemistry for MCH and Aβ plaques. MCH-positive axons from AD patients that had plaque deposition displayed the same phenotype as we observed in *App^NL-G-F^* mice – no change in the number of MCH-positive puncta but an increase in their area and a dystrophic appearance near plaques (Fig. 4d-g). This phenotype correlated with progression of the disease, as determined by the neuritic plaque score developed by The Consortium to Establish a Registry for Alzheimer’s Disease (CERAD) (Fig. 4e) or Braak neurofibrillary tangle (NFT) staging (Fig. 4f). Altogether, we find that MCH-positive axons in the hippocampal CA1 region have an increased MCH-positive puncta area and a dystrophic appearance near Aβ plaques in both mouse model and human AD samples, providing a human morphological correlate of the impaired MCH neuron activity we observe in the early stages of Aβ pathology in the mouse model.

## Discussion

Our work shows that homeostatic mechanisms transiently counteract Aβ-induced hyperactivity in the hippocampal CA1 neurons but ultimately fail to stabilize neuronal activity in *App^NL-G-F^* mice at a physiological set-point ^9^. Remarkably, fluctuations in cognition of AD patients have been reported ^3^. Here we find that MCH regulates synaptic plasticity in hippocampal neurons and that the MCH-system is dysregulated in the early stages of AD, contributing to the failure of homeostatic responses to the gradual build-up of Aβ. This supports the idea that current models for acute Aβ toxicity are oversimplified and that perturbations of the cellular regulators of homeostasis play a key role in gradual Aβ-mediated toxicity processes ^49^.

Using ST we reveal potential molecular sensors and effectors that detect and correct deviations from the physiological set-point ^9,10^. The advantage of the ST approach is that it covers cellular niches in the brain, yielding the transcriptional profiles of CA1 pyramidal neurons but also of the surrounding environment ^23^. This was instrumental to identify *Pmch* mRNA changes in the CA1 region that arise from alterations in projecting MCH-axons from the LHA. The interaction of the hippocampus with LHA across the awake-sleep cycle has been implicated in memory consolidation ^50^ and persistent wakefulness increases the risk for seizures ^51,52^ due to impaired downscaling of synapses. Our observations show that MCH modulates hippocampal synaptic plasticity and suggest that the MCH-system is involved in safeguarding neurons from hyperexcitability by downscaling synapses. Thus, the failure of the MCH system in the early stages of AD provides a cellular and molecular underpinning of previously observed aberrant neuronal activity, sleep disturbances and impaired memory formation in AD. Remarkably, increased MCH levels in the cerebral spinal fluid of mild-cognitive impaired patients have been reported ^53^. Therapeutically targeting the MCH-system to modulate neuronal homeostasis may therefore have major implications for the treatment of early stages of AD.

## Acknowledgements

We thank Takaomi Saito for the generation of the transgenic mice, Antoine Adamantidis for critical reading and experimental advice, Pierre Vanderhaeghen, Patrik Verstreken, Lucas Baltussen, Anna Martinez, Renzo Mancuso, Nuno Apóstolo and Luis Ribeiro for critical reading of the manuscript, and De Wit and De Strooper lab members for helpful discussion and comments. We thank the VIB-KU Leuven BioImaging core and Nucleomics core for experimental help. Leica SP8x confocal microscope was provided by InfraMouse (KU Leuven-VIB) through a Hercules type 3 project (ZW09-03). S.C. is supported by Fonds voor Wetenschappelijk Onderzoek (FWO, Belgium), Stichting Alzheimer Onderzoek (SAO, Belgium), the Alzheimer’s Association (USA); DRT is supported by FWO grants G0F8516N, G065721N and SAO-FRA grant 2020/017. J.d.W is supported by SAO Grant 2019/0013, FWO Odysseus Grant, FWO EOS Grant G0H2818N, and Methusalem Grant of KU Leuven/Flemish Government; B.d.S is supported by the European Research Council ERC-CELLPHASE_AD834682 (EU), Methusalem Grant of KU Leuven/Flemish Government, Geneeskundige Stichting Koningin Elisabeth (Belgium), and Bax-Vanluffelen (Belgium).

## Author Contributions

S.C., J.d.W and B.d.S. conceived the study and designed experiments. S.C. performed patch-clamp experiments. E.C. performed MEAs experiments. S.C., G.O., J.v.L and L.v.B. performed immunohistochemistry, microscopy imaging and analysis. N.T, S.P, K.D, A.L., M.F and W.C. were involved in RNAsequencing and transcriptomics analysis. S.C. performed western blots. K.C. performed elisa assays. D.R.T. performed pathology scoring on human samples. J.v.B. maintained animal colonies and genotyped animals. J.v.B., S.C. and G.O. prepared animal brain samples. S.C., J.d.W and B.d.S. wrote the manuscript.

