## Supplemental Figures for "Melanin Concentrating Hormone- and sleep-dependent synaptic downscaling is impaired in Alzheimer’s Disease"

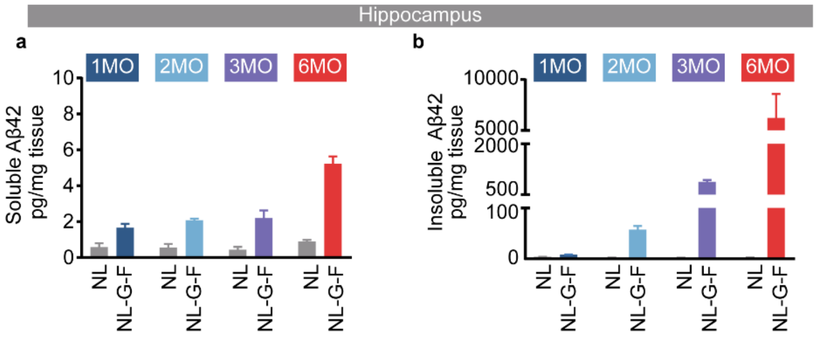


**Extended Data Fig.1**

**a,b,** Quantification of **(a)** soluble and **(b)** insoluble A$\beta_{42}$ using meso ELISA on *App*$^{NL}$ and *App*$^{NL\text{-}G\text{-}F}$ mice hippocampal lysates at 1, 2, 3 and 6 months (MO). Number of animals: n=3 animals per each time point and genotype. *App*$^{NL}$ mouse was used as control as it contains the human *App* gene but with only one mutation in (NL) and does not develop Aβ plaques when compared with *App*$^{NL\text{-}G\text{-}F}$

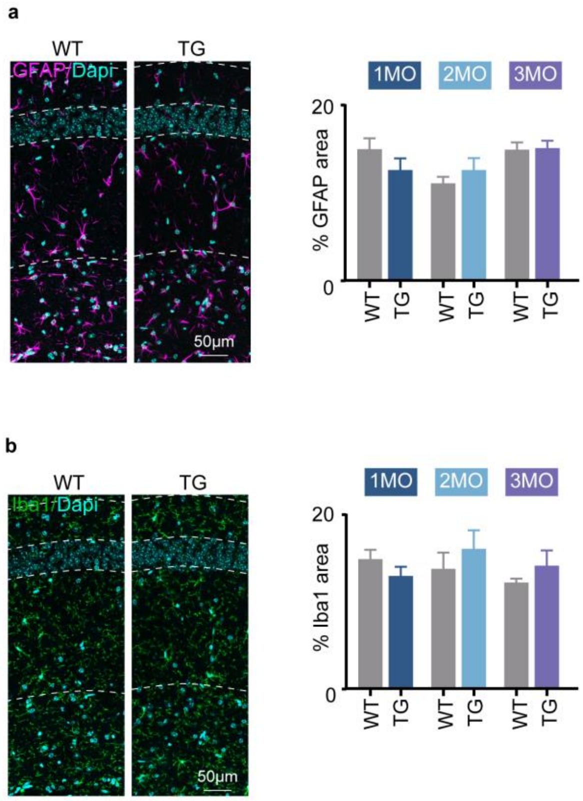


**Extended Data Fig.2**

**a,b,** Representative image and area covered by signal on CA1 hippocampal sections from WT and *App*$^{\text{NL-G-F}}$ (TG) mice at 1, 2 and 3 months (MO) immunostained for **(a)** astrocyte-marker GFAP (magenta) and **(b)** marker of microglia activation Iba1 (green). Nuclei are labeled with DAPI (cyan).

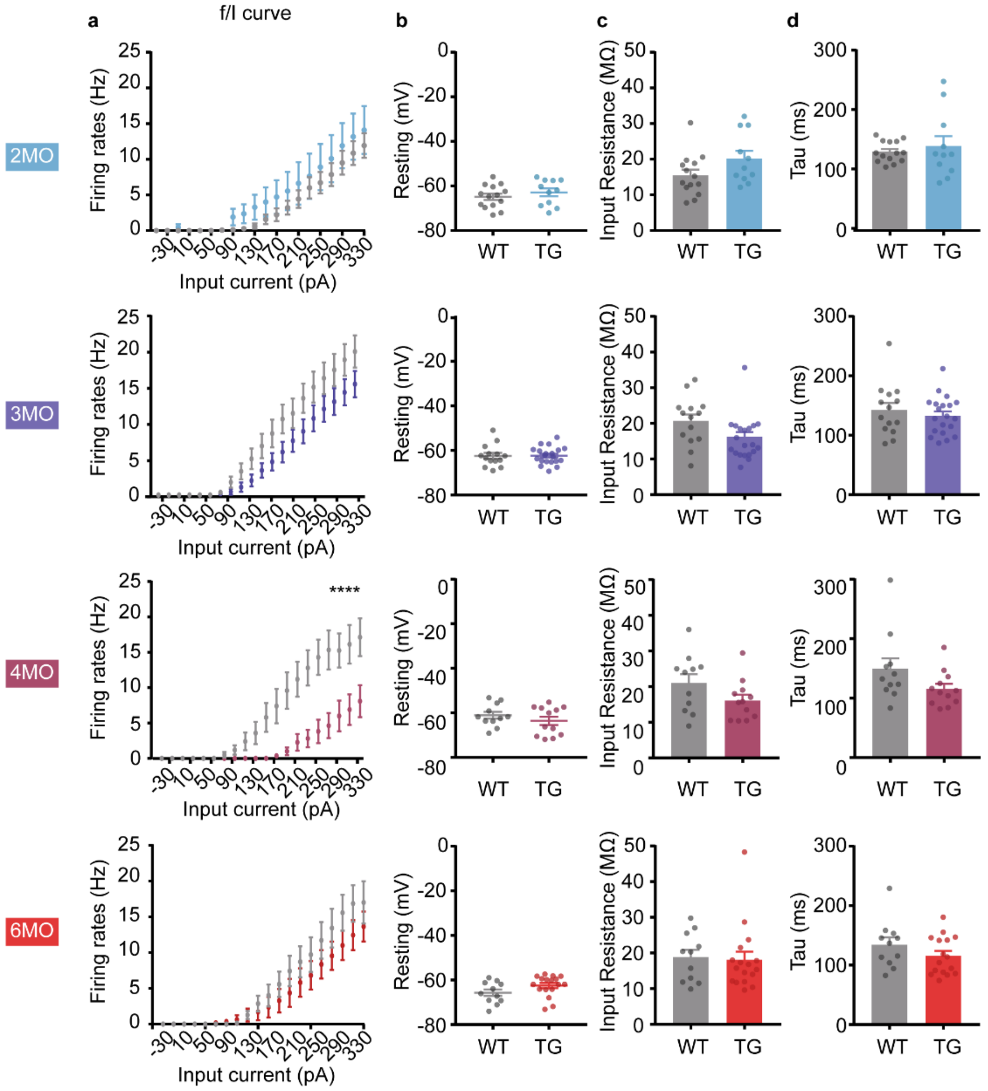


**Extended Data Fig.3**

**a,b,c,d** Whole-cell current clamp recordings of intrinsic properties from CA1 pyramidal neurons from WT and *App*$^{NL-G-F}$ (TG) at different months (MO). **(a)** intrinsic excitability, firing rate in function of somatic current injection, **(b)** resting membrane potential, **(c)** input resistance and **(d)** Tau membrane constant. Number of neurons, n=14, WT and n=20, TG 3MO; n=11, WT and n=12, TG 4MO, n=11, WT and n=17, TG 6MO. n=3 animals for each genotype. **(a)** ANOVA test for multiple comparisons, ****P < 0.0001. **(b,c,d)** Two-tailed unpaired *t*-test.

a

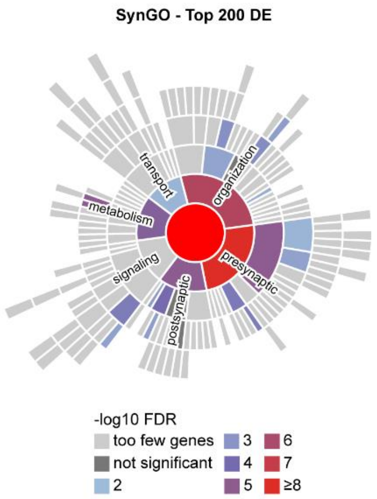


**Extended Data Fig.4**

**a,** Sunburst visualization of Synaptic Gene Ontology (SynGO) enriched ontology terms on the top 200 DE genes (based on P value), using Fisher's Exact Test, coloured by -log10 FDR (False Discovery Rate).

a

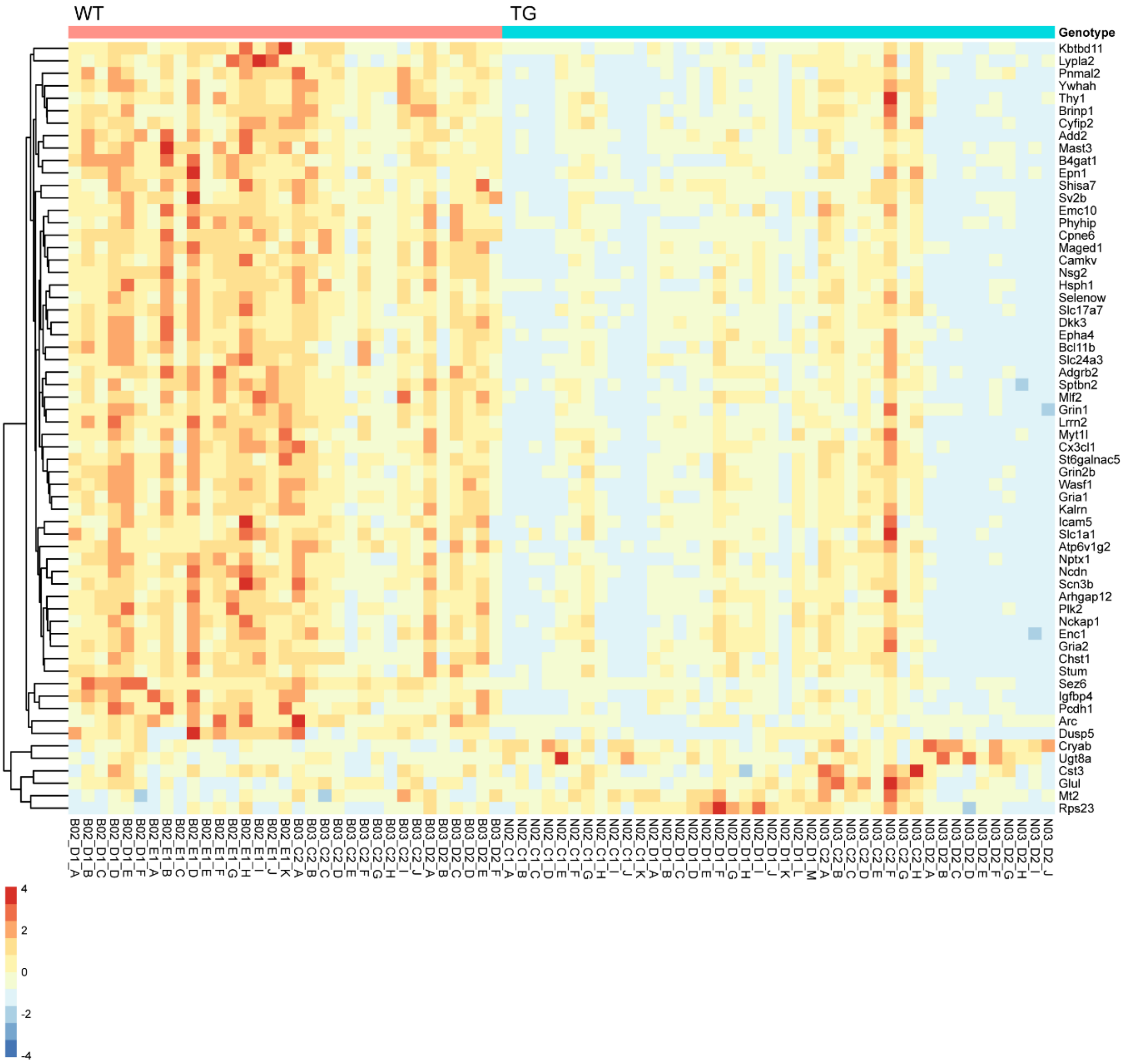


### Extended Data Fig.5

**a,** Heatmap showing scaled expression of DE genes (in ST) that appear in at least one homeostatic plasticity datasets [1–4]. Columns represent spots taken from the CA1 region of the ST data, from multiple tissue slides of 2 *App*$^{NL-G-F}$ (TG) mice and 2 wildtype (WT) mice at 3.5 months (mouse genotypes are shown as coloured bars across the top). Mouse information metadata is available in Supplementary Table 4

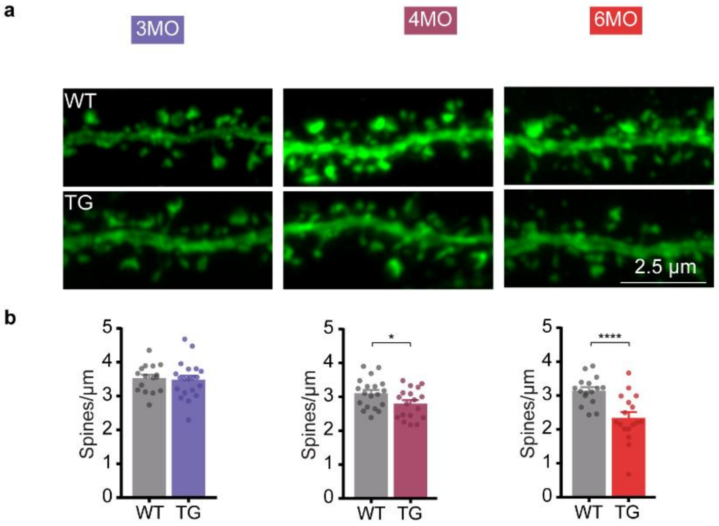


**Extended Data Fig.6**

**a,b,** Spine analysis of CA1 pyramidal neuron proximal dendrites labelled with GFP from WT and *App*$^{NL\text{-}G\text{-}F}$ (TG) at different months (MO). **(a)** representative images and **(b)** quantification of spine number per dendrite length. n=3 animals per time point and genotype. Two-tailed unpaired *t*-test (*P < 0.05, ****P < 0.0001).

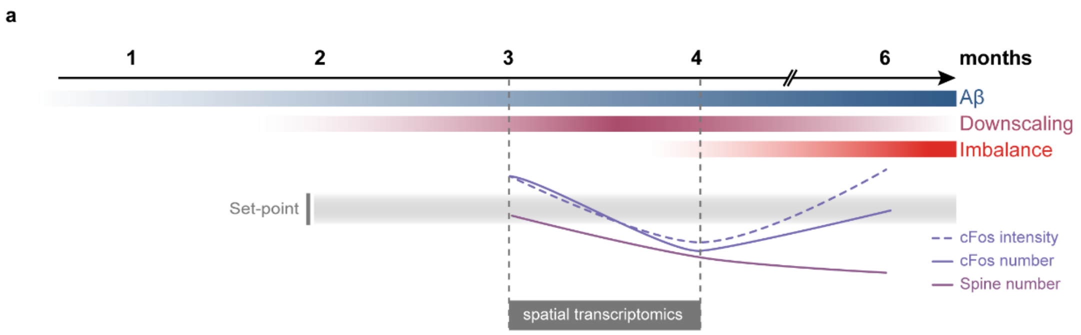


**Extended Data Fig.7**

**a,** Schematic summary of data shown in Fig. 1a-c and Extended Data Fig. 3 and 6

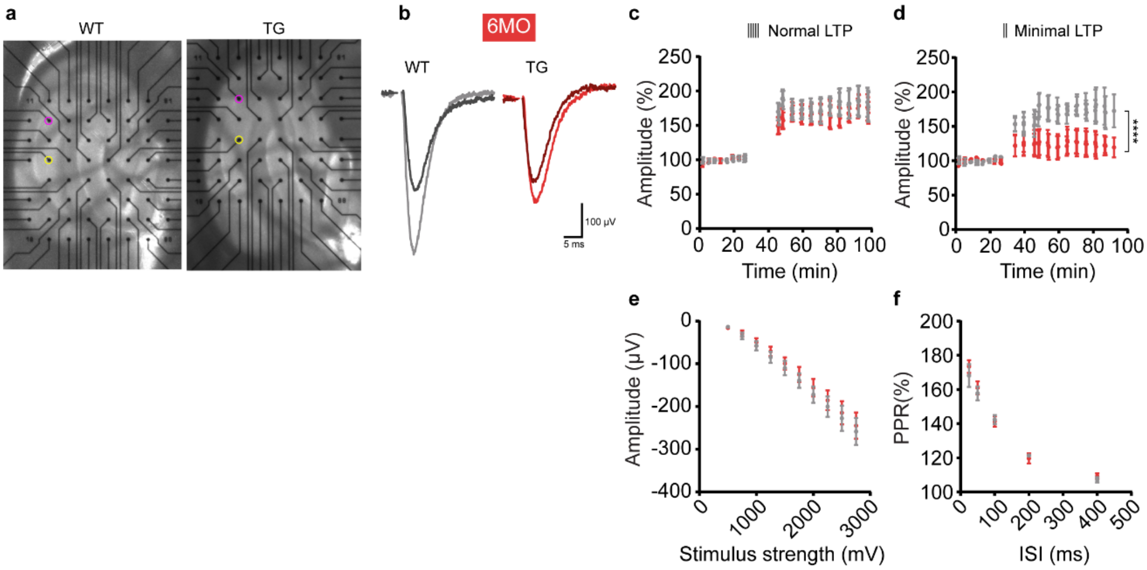


**Extended Data Fig.8**

**a,** Images of control (WT, left) and *App*$^{NL\text{-}G\text{-}F}$ (TG, right) acute hippocampal slices on the multielectrode array (MEA2100, Multichannel Systems) used for field excitatory post-synaptic potential (fEPSP) recordings. Stimulation and recording electrodes are indicated with purple and yellow circles, respectively. **b,** Example traces of fEPSP responses in WT and TG before (dark traces) and 55 minutes after minimal LTP induction (light traces). **c,** LTP induced in CA1 region by Schaffer colateral (SC) pathway stimulation using 3 theta burst stimulations (100 stimuli each). **d,** LTP induced in CA1 region using minimized theta burst stimulation paradigm (2 bursts of 100 Hz, 75 stimuli each). **e,** Input-output (IO) relationship for WT and TG slices. **f**, Paired-pulse ration (PPR) stimulations of SC pathway. Stimulation intensities used for LTP and paired pulse ratio recordings were calculated from CA1 region using SC stimulation in WT and TG slices (inter stimulation intervals: 25, 50, 100, 200 and 400 ms).

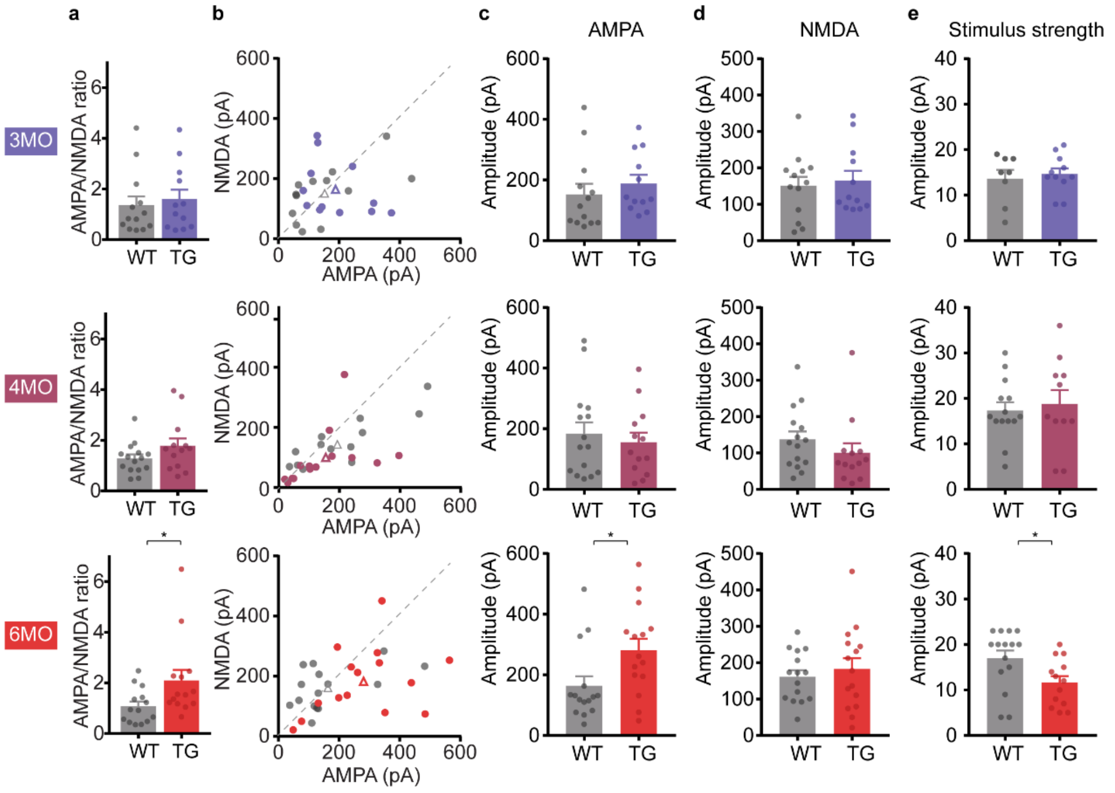


**Extended Data Fig.9**

**a,b,c,d,e** Whole-cell voltage clamp recordings of AMPA (-70mV) and NMDA (+40 mV) currents from CA1 pyramidal neurons from WT and *App*$^{NL\text{-}G\text{-}F}$ (TG) at different months (MO) **(a)** AMPA/NMDA ratio, **(b,c,d)** AMPA and NMDA raw current amplitudes, **(e)** electrode stimulation strength. Number of neurons, n=13, WT and n=12, TG 3MO; n=15, WT and n=13, TG 4MO, n=15, WT and n=15, TG 6MO. n=3 animals for each genotype. Two-tailed unpaired *t*-test (*P < 0.05).

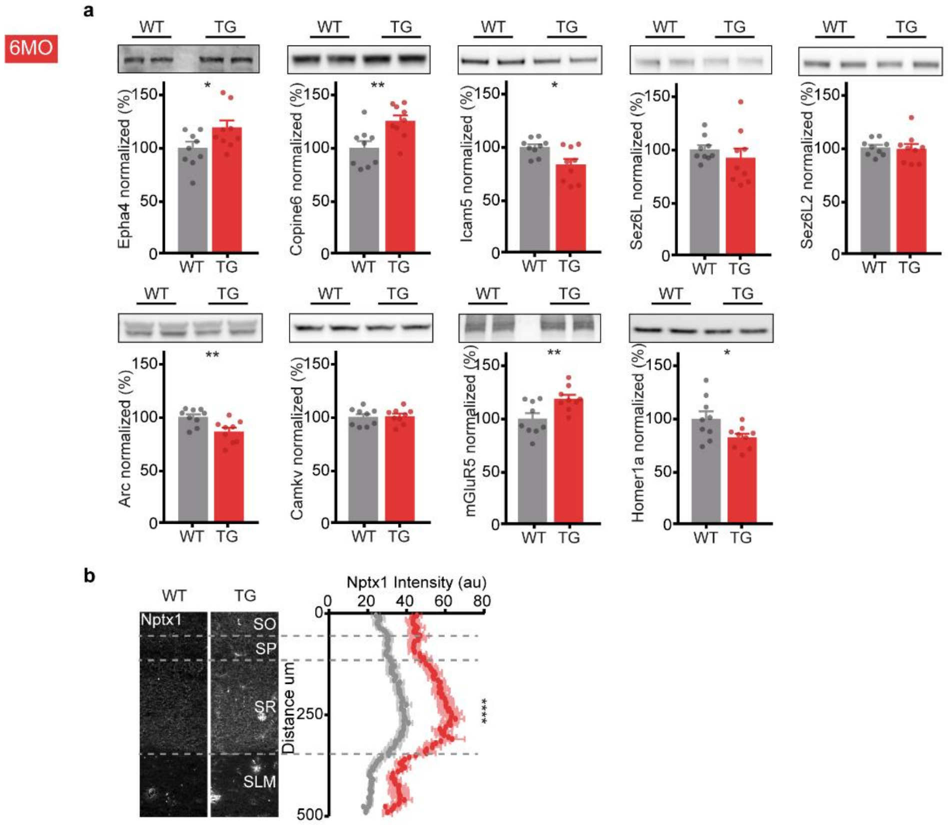


**Extended Data Fig.10**

**a,** Hippocampal synaptosomes from WT and *App*$^{NL\text{-}G\text{-}F}$ (TG) at 6 months were loaded on a western blot and protein levels of several DE genes from the ST analysis were assessed using immunolabeling with different antibodies. Representative western blots and quantifications showing normalized band intensities to WT are shown for each antibody. n=9 WT; n=9 TG. Two-tailed unpaired *t*-test (*P < 0.05, **P < 0.01).

**b,** Brain coronal sections from WT and *App*$^{NL\text{-}G\text{-}F}$ (TG) at 6 months were immunostained for Nptx1 protein and intensity analyzed along SO-SLM axis. ANOVA test for multiple comparisons, ****P < 0.0001. SO – stratum oriens, SP – stratum pyramidale, SR – stratum radiatum, SLM – stratum lacunosum moleculare.

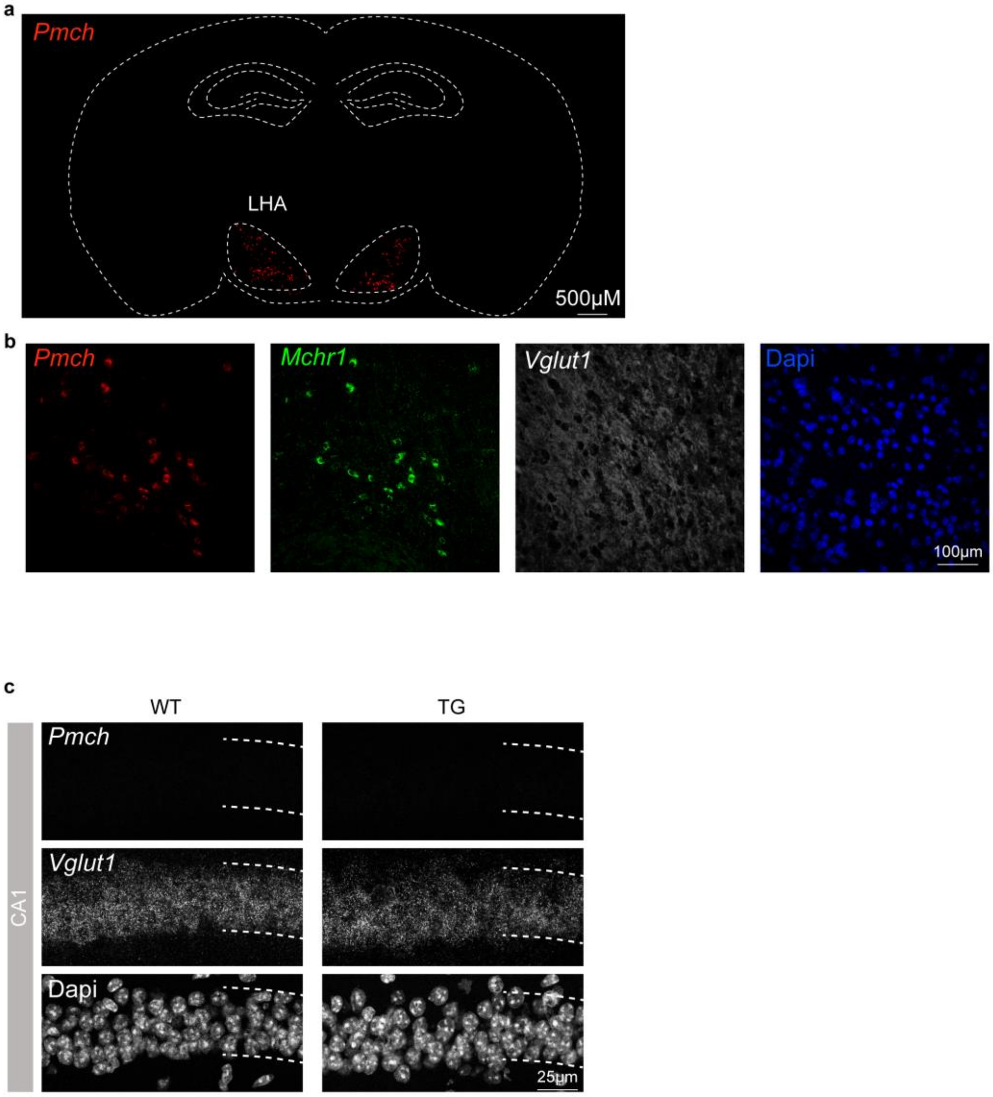


**Extended Data Fig.11**

**a,b,** RNAscope on 10μm WT mouse brain coronal section showing **(a)** *Pmch* expression in whole section and **(b)** *Pmch*, *Mchr1* and *Vglut1* expression in the lateral hypothalamic area (LHA) shown by higher magnifications images of this brain region. **(c)** *Pmch* and *Vglut1* expression in CA1 region of the hippocampus from WT and $App^{NL\text{-}G\text{-}F}$ (TG) at 3.5 months old.

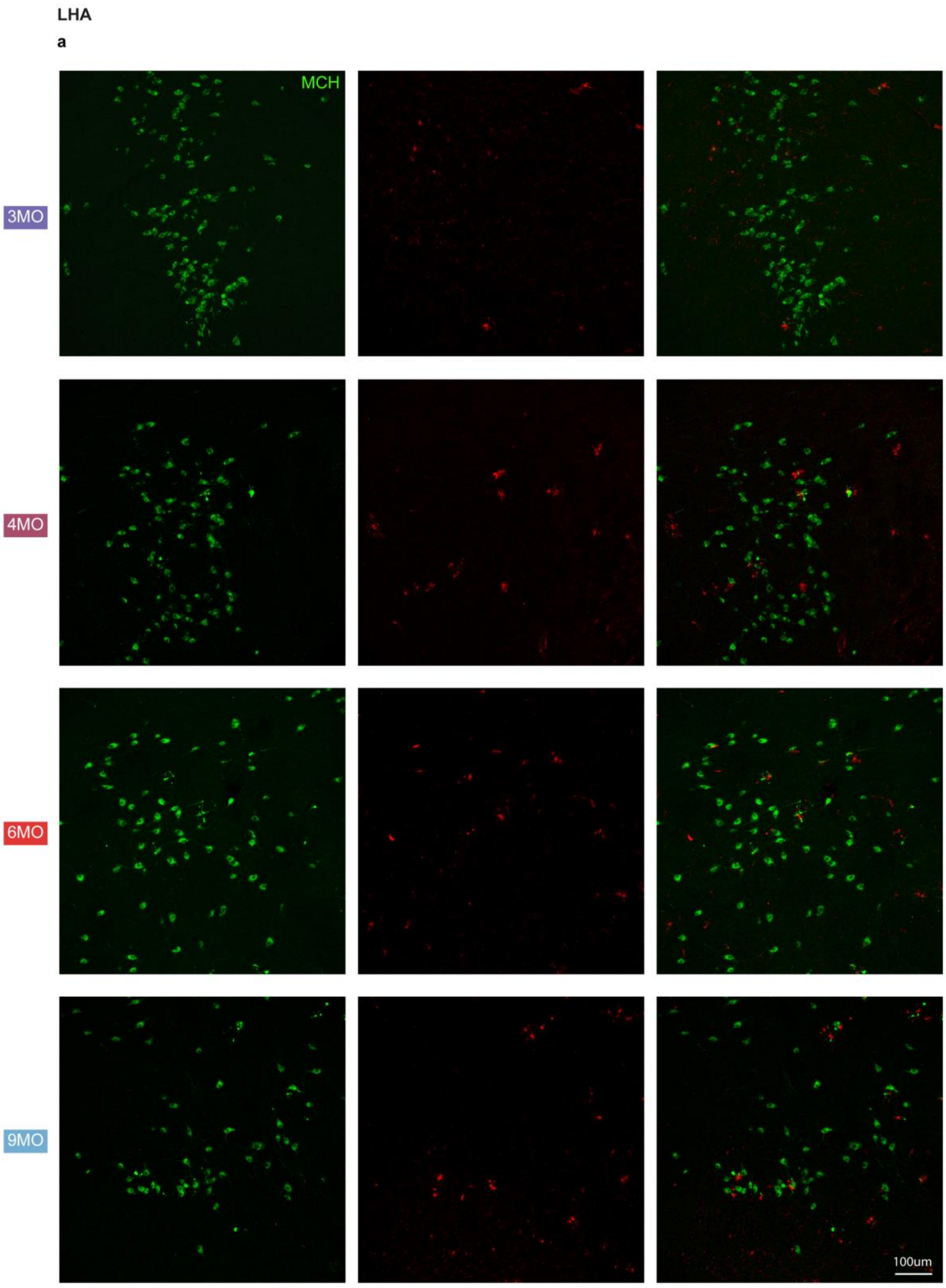


**Extended Data Fig.12**

**a,** Images from LHA region on a 50 µm coronal brain section immunostained for MCH (green) and Aβ (6E10 antibody - red) from *App*$^{NL\text{-}G\text{-}F}$ (TG) at 3, 4, 6 and 9 months (MO).

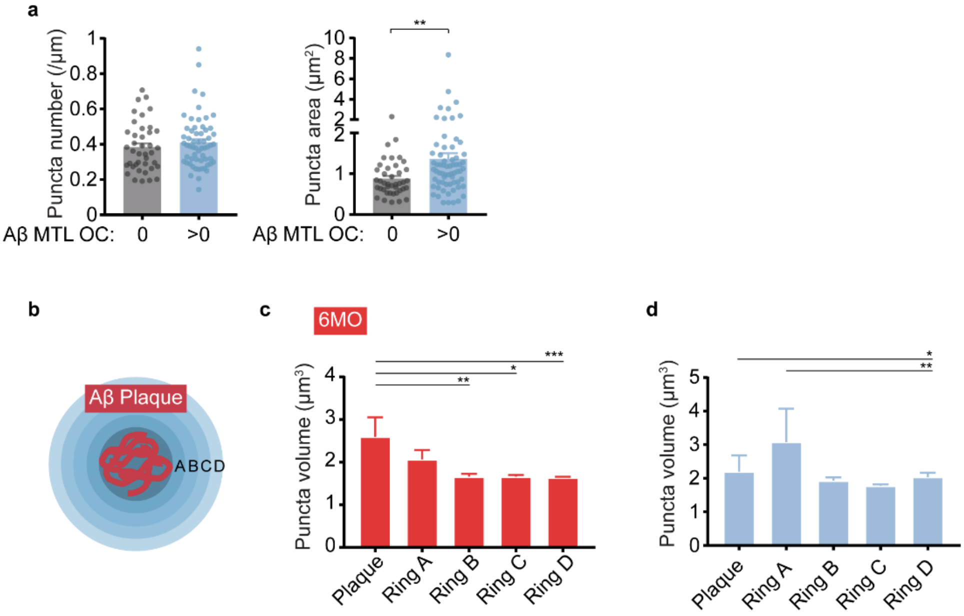


**Extended Data Fig.13**

**a,** Quantification of the number of MCH puncta along axon length and respective puncta area in relation to Aβ-MTL OC scoring in AD patients. unpaired Mann-Whitney test (**P < 0.01).

**b,c,d** Quantification of MCH puncta area nearby Aβ plaques stained by 6E10 antibody in **(c)** *App*$^{NL\text{-}G\text{-}F}$ at 6 months (MO) and **(d)** AD patients. unpaired Mann-Whitney test (*P < 0.05, **P < 0.01, ***P < 0.001).

CA1

a

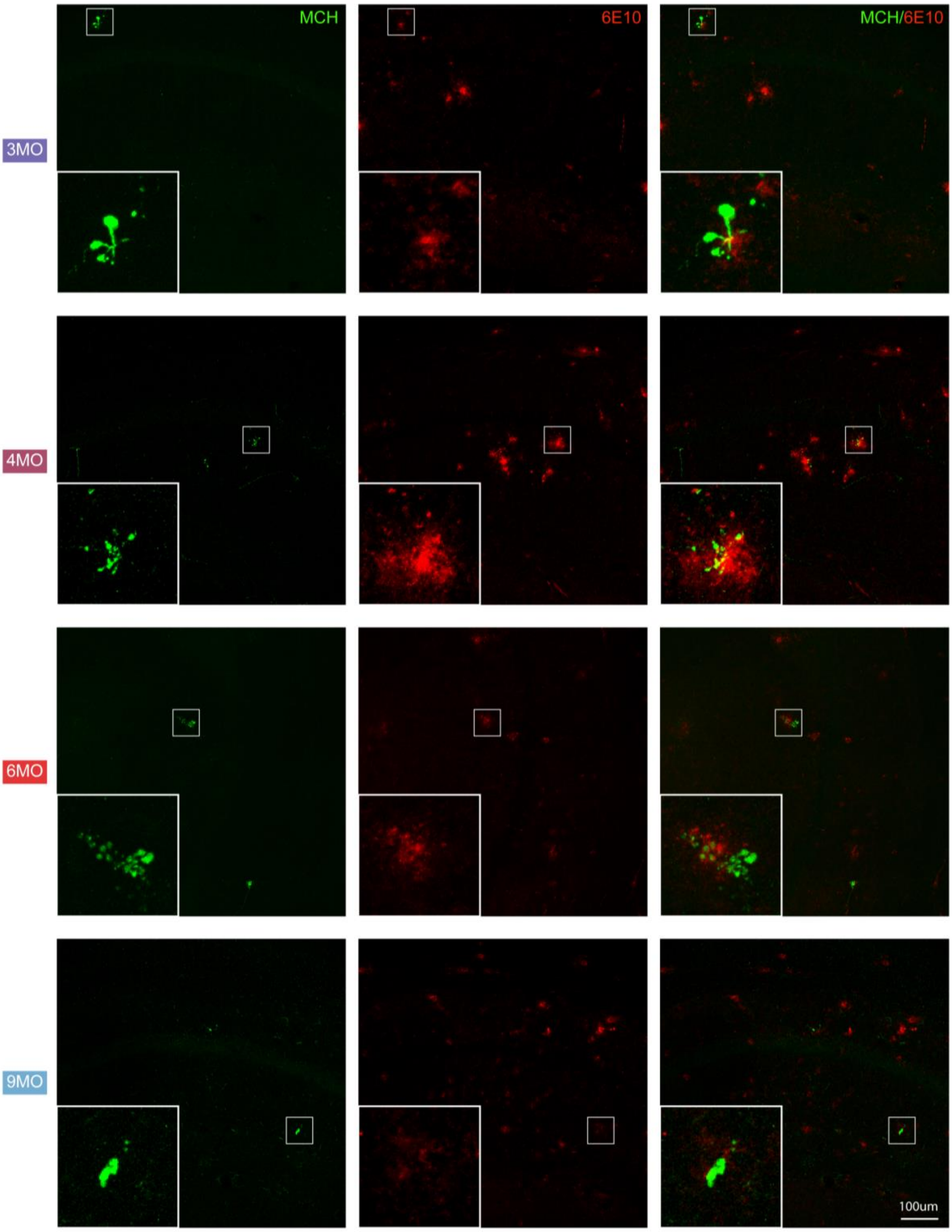


**Extended Data Fig.14**

**a,** Images from CA1 region on a 50μm coronal brain section immunostained for MCH (green) and Aβ (6E10 antibody - red) from *App*$^{NL\text{-}G\text{-}F}$ (TG) at 3, 4, 6 and 9 months (MO).
