## Supplementary material for "Melanin Concentrating Hormone- and sleep-dependent synaptic downscaling is impaired in Alzheimer’s Disease": Methods

#### Mice

All mouse lines were maintained on a C57BL/6J background, bred in-house and raised in a temperature- and humidity-controlled room with a 14-10h light–dark cycle (lights on from 7h00 to 21h00). *App*$^{NL\text{-}G\text{-}F}$ knock-in [1] mice express Swedish (KM670/671NL), Beyreuther/Iberian (I716F), and Arctic (E693G) mutations in the App gene under the endogenous promoter on the C57BL/6J background. *App*$^{NL\text{-}G\text{-}F}$ mice were backcrossed for at least 2 generations with C57BL/6J mice in the De Strooper lab. All experimental protocols were approved by institutional Animal Care and Research Advisory Committee of the KU Leuven (ECD P183/2017) and were performed in accordance with the Animal Welfare Committee guidelines of the KU Leuven, Belgium. The health and welfare of the animals was supervised by a designated veterinarian. The KU Leuven animal facilities comply with all appropriate standards (cages, space per animal, temperature, light, humidity, food, water), and all cages are enriched with materials that allow the animals to exert their natural behaviour.

#### Antibodies

Anti-mouse EphA4 (Invitrogen 37-1600, 1/500), anti-mouse Copine 6 (santacruz sc-136357, 1/500), anti-mouse Neuronal Pentraxin 1 (BD 610369, 1/2000), anti-mouse GluR1 (Milipore MAB2263, 1/500), anti-rabbit MCH (Phoenix, H-070-034, 1/500), anti-rabbit Camkv (Fisher PA5-101203, 1/1000), anti-mouse MCHR1 (RD MAB7938-SP, 1/1000), anti-rabbit GluR1 pSer845 (Milipore AB5849, 1/500), anti-rabbit Arc (SantaCruz sc-15325, 1/200), anti-rabbit mGluR5 (Merck AB5675, 1/500), anti-rabbit Homer1a (Synaptic Systems 160-030, 1/500, anti-sheep Sez6l (R&D AF4804, 1/1000), anti-sheep Sez6l2 (R&D AF4916, 1/1000), anti-goat Icam5 (RD AF1173, 1/1000), anti-chicken GFP (Aves GFP-1010, 1/2000), anti-guinea pig vGlut1 (Milipore AB5905, 1/5000), anti-mouse 6E10 (Biolegend 803003, 1/1000), anti-sheep Orexin (LSBio LS-B31, 1/1000), anti-mouse cFos (EnCore MCA-2H2, 1/1000), anti-rabbit cFos (Synaptic Systems 226-003, 1/1000), anti-rabbit Iba1 (Synaptic Systems 234-003, 1/1000), anti-guinea pig GFAP (Synaptic Systems 173-004, 1/1000)

#### Electrophysiology

Electrophysiology was performed at the VIB-KU Leuven Center for Brain and Disease Research Electrophysiology Expertise Unit. All recordings were performed on WT or *App*$^{NL\text{-}G\text{-}F}$ littermate pairs. For any parameter analysed, a minimum of 3 pairs were used. Mice were anesthetized with isofluorane and rapidly decapitated to prepare acute 300 μm-thick parasagittal brain slices on a Leica VT1200 vibratome. Slicing was performed in a sucrose based cutting solution (ACSF) that consisted of (in mM): 87 NaCl, 2.5 KCl, 1.25 $NaH_2PO_4$, 10 glucose, 25 $NaHCO_3$, 0.5 $CaCl_2$, 7 $MgCl_2$, 75 sucrose, 1 kynurenic acid, 5 ascorbic acid, 3 pyruvic acid (pH 7.4 with 5% $CO_2$/ 95% $O_2$). Slices were allowed to recover at 34°C for 30 min, and then maintained at room temperature in the same solution for at least 30 min before using. Pipettes were pulled on a horizontal micropipette puller (Sutter P-1000) and resistances ranged from 3 to 5 MΩ. Whole-cell voltage clamp recordings were made of CA1 pyramidal neurons at the distal region of pyramidal layer and data collected with pCLAMP 10. All recordings were done at 34°C. Input resistance, pipette series resistance and membrane holding current were monitored throughout all recordings to ensure stability and quality. Currents were sampled at 20 kHz and stored after 3 kHz low-pass Bessel filtering (Molecular Devices DigiData 1440A and Multiclamp 700B). Before analysis the data were low-pass filtered at 1 kHz. For sEPSC and evoked recordings, slices were perfused at 1-2 mL/min with ACSF consisting of (in mM): 119 NaCl, 2.5 KCl, 1 $NaH_2PO_4$, 11 glucose, 26 $NaHCO_3$, 4 $MgCl_2$, 4 $CaCl_2$ and 0.1 picrotoxin,bubbled continuously with 95% $O_2$ and 5% $CO_2$ using a Cs-methanesulfonate-based internal solution (in mM): 115 CsMSF, 20 CsCl, 10 HEPES, 2.5 $MgCl_2$, 4 ATP disodium salt, 0.4 GTP sodium salt, 10 creatine phosphate and 0.6 EGTA, adjusted to pH 7.5 and 295 mOsm. For sEPSC recordings, membrane potential was clamped at −70mV. For evoked recordings a 2-contact cluster microelectrode (FHC, CE2C55) was placed in SR of CA1 stimulate Schaffer collaterals synaptic responses over 10-15 sweeps. AMPAR-mediated EPSCs were recorded at a holding potential of −60 mV and compound AMPAR- and NMDAR-mediated EPSCs were recorded at +40 mV (NMDA component was quantified 50 ms after initiation of the combined AMPAR-and NMDAR-mediated EPSCs). Stimulus intensity was adjusted to standardize the AMPAR EPSC recorded. For intrinsic properties we perfused at 1-2 mL/min with ACSF consisting of (in mM): 124 NaCl, 2.5 KCl, 1.2 $NaH_2PO_4$, 24 $NaHCO_3$, 5 HEPES, 12.5 Glucose, 2 $MgSO_4 \cdot 7H_2O$, 2 $CaCl_2 \cdot 2H_2O$ and 0.1 picrotoxin bubbled continuously with 95% $O_2$ and 5% $CO_2$ and used a K-gluconate based internal solution (in mM): 135 KGluconate, 4 KCl, 2 NaCl, 10 HEPES, 4 EGTA, 4 Mg ATP, 0.3 Na GTP, adjusted to pH 7.25 and 295 mOsm. Somatic current injections of 20 pA steps starting at −50 until 530 pA were used for action potential profiling. sEPSCs were analysed using the Mini Analysis program (Synaptosoft). Evoked responses and intrinsic properties were quantified using Clampfit 10.7 (Axon Instruments).

**Multi-electrode array electrophysiology**

Whole brain parasagittal sections were prepared as described above. Age matched WT and $App^{NL\text{-}G\text{-}F}$ mice (~ 6 months) were used for LTP experiments. For recordings, slices were placed onto a multielectrode array (MEA 2100, Multichannel Systems) and continuously perfused with 34°C artificial cerebrospinal fluid (aCSF) solution (119 mM NaCl, 2.5 mM KCl, 1 mM NaH2PO4, 11 mM glucose, 26 mM NaHCO3, 4 mM MgCl2 and 4 mM CaCl2) at pH 7.4 and 5% CO2/ 95% O2. Field excitatory post-synaptic potentials (fEPSPs) were recorded from Schaffer collateral-CA1 synapses by stimulating and recording from the appropriate (visually identified) electrodes. Input-output curves were recorded for each slice by applying single-stimuli ranging from 500 to 2750 mV with 250 mV increments. Stimulus strength that corresponds to 35% of maximal response in the input-output curve was used for following recordings. Paired-pulse facilitation experiments were performed by applying paired stimuli with 25, 50, 100, 200 and 400 ms inter-stimulus intervals. For long-term potentiation (LTP) experiments, stable fEPSPs were recorded for 30 minutes to establish a baseline. For normal LTP induction, we applied three high frecuency trains (100 stimuli; 100 Hz) with 5 minute intervals. For minimal LTP induction, we used two shorter trains (75 stimuli; 100 Hz) with a 5 minute interval. Subsequently, post-LTP fEPSPs are measured every 5 minutes (average of three consecutive stimulations (15 seconds apart) for 55 minutes. Recordings were analyzed and processed using Multi Channel Experimenter software (Multichannel Systems).

**Primary hippocampal neuronal cultures**

Hippocampal neurons were cultured from E18 C57/Bl6 wild type mice and 1 million neurons were plated on each well of a 6-multiwell plate coated with poly-D-lysine (Millipore) and laminin (Invitrogen) coated. Neurons were maintained in Neurobasal medium (Invitrogen) supplemented with B27 and insulin (Invitrogen). Neurons were treated with $H_2O$ or 1 µM MCH peptide (Phoenix H-070-47) and collected for immunoblotting or RNA extraction on DIV12-14.

**Spatial transcriptomics**

The gene expression data obtained from ST is derived from a dataset available in the host laboratory (Chen, 2020). Raw data is available at GSE152506. Briefly, 10 µm thick coronal brain cryosections (bregma: -2.0 to -2.2) were obtained from 3.5 month old WT or $App^{NL\text{-}G\text{-}F}$ mice, layered onto a spatially barcoded array of 1007 TDs (diameter of 100 µm and a center-to-center distance of 200 µm, over an area of 6.2 mm by 6.6 mm) to collect in situ 2D-RNaseq (Lot#10001, Spatial Transcriptomics, Stockholm, Sweden). Each spot contains approximately

200 million barcoded reverse-transcription oligo(dT) primers allowing to get a global transcriptomic profile of a TD with a volume of 0.00008mm3(pr2h with r = 50 μmand h = 10 μm). Images were acquired by Zeiss Axio Scan.Z1 slidescanner (Carl Zeiss AG, Oberkochen, Germany) and library preparation was performed after imaging following the Library Preparation Manual (Spatial Transcriptomics, Stockholm, Sweden) as previously described [2]. We extracted expression data assigned to the hippocampal region of WT and *App*$^{NL\text{-}G\text{-}F}$ mice at 3.5 months of age for analysis.

*UMAP embeddings.* We used Seurat v3.1.4 [3] to cluster hippocampal ST data as follows: normalized expression data (counts per million normalized to library size and log-transformed) was used as input. We scaled the expression data based on the 2,000 most variable genes as calculated using Seurat's variance stabilizing transform algorithm, regressing on number of reads at the same time. A principal components analysis was run, and used the top 10 principal components to calculate nearest neighbors and UMAP coordinates, leaving all other Seurat parameters as default.

*Differential expression and enrichment.* Differential expression of the CA1 pyramidal layer (sp) - comparing WT or *App*$^{NL\text{-}G\text{-}F}$ genotypes - was performed as described in the original publication. Briefly, generalized linear models were fit, and differential expression was tested using EdgeR's quasi-likelihood F test [4]. We defined differentially expressed (DE) genes as those genes with False Discovery Rate (FDR) < 0.05 as significant. The full results of the DE are available in Supplementary table 1.

*Gene Onthology (GO).* To test for enrichment of ST DE genes, we extracted the top 200 genes (according to P value) and submitted to Gorilla [5], using default parameters. GO categories were sorted by P value and top 8 ordered by normalized enrichment score (Fisher's Exact Test). 5 most enriched GO categories are shown in the bar plot. The genes annotated in each GO term are available in Supplementary Table 2.

*Annotations in SynGO*. We extracted the ST DE top 200 genes (according to P value) and submitted SynGO v1.0 database using default parameters (after using the SynGO conversion tool to convert mouse gene IDs to human IDs) [6]. SynGO genes are available in Supplementary Table 3.

*Comparisons with homeostatic plasticity datasets*. We extracted homeostatic synaptic plasticity data from the following studies: 1) RNA-Seq data from a study of the transcriptional program responsible for synaptic upscaling during activity suppression (all genes affected by TTX or Bicuculin [7], a characterization of the surfaceome of primary neuronal cultures (all total proteins and surface proteins changing with TTX or Bicuculin treatment, $p < 0.05$, Supplementary table 3) [8]; 3) a study of the synaptic proteome of the primary sensory cortex (proteins up- and down-regulated in sensory-deprived cortex [9]; 4) a single-cell RNA-Seq study of activity-dependent transcriptional changes in mouse excitatory neurons[10] (genes changing in clusters ExcL clusters in Supplementary Table 3). Protein names / IDs from proteomics data were converted to gene names using the UniProt mapping tool (https://www.uniprot.org/uploadlists/), or in the case where a match could not be found, manual annotation. This comparison is available in Supplementary Table 3.

*Heatmaps.* Heatmaps were generated using log-normalized expression data (normalized using EgdeR's *cpm* function), then scaled per row. IDs from the original ST data have been modified for visualization purposes.

**Bulk RNAsequencing**

For bulk RNA-seq, two independent cultures of hippocampal neurons were used. In each culture neurons, we treated 2 wells with $H_20$ (veihcle) and 2 wells with 1 µM MCH peptide (Phoenix H-070-47). RNA was extracted using RNeasy Mini Kit (Qiagen 74104). RNA purity (260/280 and 260/230 ratios) and integrity were assessed using Nanodrop ND-1000 (Nanodrop Technologies, Wilmington, DE, USA) and Agilent 2100 Bioanalyzer with High Sensitivity chips (Agilent Technologies, Inc., Santa Clara, CA, USA) and Qubit 3.0 Fluorometer (Life Technologies, Carlsbad, CA, USA), respectively. RNA integrity values of the samples ranged from 7.9 to 9.3 (median = 8.6). Library preparation from total RNA extract and sequencing were performed at the VIB Nucleomics Core (Leuven, Belgium). Briefly, 1ug of total RNA extract per sample was enriched for mRNA molecules using poly-T oligo attached magnetic beads. The enriched poly-A mRNA species is subjected to fragmentation and reverse transcriptions using random primers and the subsequent library processing was done using standard Illumina TruSeq Stranded mRNA Sample Prep Kit (protocol version 15031047 Rev.E). Libraries were sequenced using an Illumina NovaSeq 6000 instrument (Illumina, San Diego, CA, USA) at an average depth of approximately 31 million reads. Raw reads were pre-processed, mapped and quantified against the GRCm38 Mus musculus genome using the nf-core/rnaseq pipeline (v3.0, https://doi.org/10.5281/zenodo.1400710).

*Differential analysis:* Salmon quantifications from the nf-core/rnaseq pipeline were imported into R v4.0.3 using the tximport library. Resulting counts were normalised and differential analysis was performed using DEseq2 (v.1.13.0). Genes were tested for differential expression between H2O treated and MCH treated samples and those with an adjusted p-value (Benjamini & Hochberg) <0.05 were deemed significant.

*Functional analysis*. To test for enrichment of differentially expressed genes, the top 200 genes (according to p value) were submitted to Gorilla [5] to test for enrichment, using default parameters.

**Immunohistochemistry, imaging and quantification**

Mice were anesthetized with ketamine (0.2mg/g body weight) and xylazine (0,02mg/g body weight) intraperitoneally administered, perfused with 1x PBS for 1 min followed by 10 min of 4% paraformaldehyde (PFA) in PBS. Brains were post-fixed for 4h in 4%PFA, washed in 1xPBS and embeded in 3% agarose. 50 µm sections were prepared in a vibratome (Campden instruments 7000smz). Sections were permeabilized with 0.5% triton in PBS-0.2% gelatin for 30 minutes, blocked for 2h in 10% normal horse serum and 0.5% triton in PBS-0.2% gelatin. Primary antibodies were incubated for 48h and secondary antibodies for 24h. Primary and secondary antibodies were diluted in 5% normal horse serum and 0.5% triton in PBS-0.2% gelatin. Hoechst was used as a nuclear stain (5nM in PBS).

*SO-SLM axis intensity.* Confocal images were taken on a Leica TCS SP8 at 20x magnification. Intensity measurements (NPTX1 images) were performed in Fiji by measuring a plot profile of the selected CA1 region (same ROI template used for all images) and intensity values across the same distance point for all images were averaged.

*cFos CA1 region.* Confocal images were taken on a Leica TCS SP8 at 63x magnification. Analysis was performed with IMARIS 9.5.1. Briefly, an ROI was defined around CA1 pyramidal layer. Next, the total number of cells was taken based on Hoechst signal. For cFos signal, a surface tool was used to create a 3-dimensional surface according to the signal. The percentage of cFos positive cells was taken based on the ratio of Hoechst positive and cFos positive cells. For cFos intensity, mean values of intensity was taken per each cFos positive cell surface.

*cFos LHA region.* Confocal images were taken on a Leica TCS SP8 at 20x magnification. Analysis was performed with IMARIS 9.5.1. Total number of MCH or Orexin positive cells was counted manually. cFos positive cells were selected using the spot tool. The ratio of MCH or Orexin cells positive for cFos was manually determined by overlapping of the signals.

*Spine analysis.* For spine analysis, 80 µm sections were prepared and immunostained with anti-GFP and CA1 pyramidal neurons were imaged with a Zeiss LSM880 confocal microscope with an Airyscan detector. 3-4 neurons were selected per animal and 2-3 secondary dendrites were randomly selected within this ROI for analysis. Spines were quantified only from dendrites with a length of at least 20 µm. Dendritic protrusions and length were quantified in Imaris software.

*MCH morphology analysis.* MCH positive axons in all laminas of CA1 region were imaged in a LSM880, at 63x magnification and 1x zoom. Stacks of 1µm were acquired. Analysis was performed using Image J. Briefly, axon length was determined. MCH positive puncta were determined using the threshold tool and number and areas of puncta were determined using the Analyze particles tool. Particle numbers were normalized to axon length.

*MCH puncta volume analysis.* MCH axons nearby plaques, identified by 6E10 signal, were imaged in a LSM880, at 63x magnification and 1x zoom. Stacks of 0.6µm spanning the entire plaque were acquired. For quantification, NIS-elements software 5.20.01 (Nikon Instruments Europe BV.) was used. The edges of the plaque were detected with a mask and rings A to D were ROI of a segmented single nucleus (based on 6E10 staining) that were expanded by 10µm. The volume of each MCH puncta inside these rings was computed.

### Human Tissue

Brain tissues were collected in accordance with the applicable laws in Belgium and Germany. The recruitment protocols for collecting the brains received from the Municipal hospital in Offenbach/Main (Germany) were approved by the ethical committees of the University of Ulm (Germany) and of UZ-Leuven (Belgium). This study was approved by the UZ Leuven ethical committee (Belgium). Brains were fixed in a 4% aqueous solution of formaldehyde for approximately 2–4 weeks. The brain hemispheres were cut into 1 cm frontal slabs and stored in polyethylene glycol. Medial temporal lobes were dissected and embedded partially in polyethylene glycol and partially in paraffin. For neuropathological diagnosis, paraffin sections were stained with hematoxylin and eosin (H&E), the Gallyas silver method, p-tau (AT8, Pierce, 1/1000), and Aβ (4G8, Senetec, 1/5000, formic acid pretreatment). The Braak NFT-stages[11] and the phase of Aβ plaque deposition [12] were determined as recommended to determine the degree of AD pathology [13]. None of the cases included in this study showed signs of hypoxemia-related neuron damage. For immunohistochemistry, 150-200 µm PEG sections were prepared in a vibratome and transferred to 70% ethanol solution. Briefly, sections were incubated with 88% formic acid and 0.1% sodium borohydride for 20 minutes. Immunohistochemistry in human sections was performed as described above for mouse sections. Primary and secondary antibodies were incubated for 48 hours.

### AAV production

HEK293T were seeded in DMEM (Invitrogen) containing 10% FBS (Invitrogen). Transfection mix, containing PEI and OptiMEM (Invitrogen) and adenovirus helper plasmid (pAdΔF6), a packaging plasmid pAAV2/1 rep-cap 2,1 (Pennvector Core PL-T-PV0001) and pCAG-GFP plasmid (addgene 11150), were added to the cells in DMEM containing 1%FBS. Cells were incubated in DMEM containing 5%FBS for 3 days. Cells were harvested, centrifuged at

1000 × g at 4 °C for 10 min and pellets were lysed in lysis buffer (150 mM NaCl and 50 mM Tris HCl-pH 8.5). Lysates were further frozen and thawed for 3 times, centrifuged at 2000 × g at 4 °C for 5 min and Benzonase (Sigma) added at a concentration of 50 U/ml to supernatants for 30 min at 37 °C. Lysates were centrifuged at 5000 × g for 20 min at RT. OptiPrep iodixanol (Sigma) gradients of 15%, 25%, 40% and 60% were prepared with 5 M NaCl, 5× PBS with 1 mM MgCl2 and 2.5 mM KCl (5× PBS-MK) and sterile H2O, and layered in 25 × 77 mm OptiSeal tubes (Beckman Coulter). Supernatants loaded on top of gradients and centrifuged at 300,000 × g and 12 °C for 100 min in the Optima XE-100 Ultracentrifuge (Beckman Coulter). Next, AAVs were collected with an 18 G needle (Beckman Coulter) from between the 40 and 60% layers, and diluted in 5 ml 1× PBS-MK. AAVs were desalted and concentrated by centrifugation at 5000 × g for 30 min at 20 °C in a prerinsed Amicon Ultra-15 filter (Millipore) in 1× PBS-MK., aliquoted and stored at −80 °C. Purity was assessed by SDS-PAGE and silver staining.

**Stereotactic injection**

Mice were intraperitoneally injected with buprenorphine 0.05 mg/kg body weight (Vetergesic), anesthetized with 5% isoflurane and Duratears applied to the eyes. Mice were placed in a mouse stereotact frame equipped with gas anesthesia head holder (KOPF). During the rest of the procedure 2.5% isoflurane was constantly administered. After shaving and disinfecting the mouse's head, local anesthesia was administered by a subcutaneous injection with 100 µl lidocaine (xylocain 1%). After 5 min an incision was made in the skin. AAVs were injected using a glass pipette pulled on a Sutter P-1000 and placed on a Nanoject III (Drummond) for loading control. The pipette was slowly lowered to the target site and remained in place for 1 min. 50 nl of virus was injected at 1nl/sec and the pipette was removed 5 min after infusion was complete. Injections were targeted to the CA1 (anterior-posterior: −2.4 mm from bregma, medial-lateral:+-2.0 mm from sagittal suture, − 1.5 mm dorsal–ventral relative to surface of the skull). After capillary removal, the burr hole was filled with bone wax (Fine science tools 19009-00). The skin was closed using veterinary tissue adhesive (Dermafuse). Post-surgery analgesia buprenorphine (0.1mg/kg body weight) was administered 4-6h after surgery. Mice were kept for 30 days before perfusion and spine analysis.

**RNAscope**

Brains were freshly dissected and frozen in OCT compound and isopentane. 10 µm sections were prepared on a cryostat (Leica) and fixed in 4%PFA for 10 min. RNAscope hybridization was performed using the RNAscope Multiplex Fluorescent Reagent Kit v2 Assay (Advanced Cell Diagnostics). After 4 × 10 min PBS washes, tissue sections were treated with pretreatment solutions and then incubated with RNAscope probes (*Mchr1*, *Pmch*, *vGlut1*), followed by amplifying hybridization processes. DAPI was used as a nuclear stain. Prolong Gold Antifade (Thermo Scientific) was used to mount slides. Confocal images were taken on a Leica TCS SP8 microscope. Tile scans were taken on Axio Scan.Z1 slide scanner (Zeiss).

**Synaptosomal preparations**

Hippocampi from WT or $App^{NL\text{-}G\text{-}F}$ mice that were 6 months of age were dissected quickly in ice-cold Hank's Balanced Salt Solution (HBSS) and homogenized using a Dounce homogenizer in homogenization buffer (0.32 M sucrose, 5 mM Trizma Base, 1 mM MgCl2, pH 7.4) with protease inhibitors. Homogenate was spun at 1000 × g to pellet nuclei and large cell debris. Supernatant was spun at 14,000 × g for 20 min at 4 °C to pellet synaptosomes (P2). P2 was resuspend in 1% Triton lysis buffer (50 mM Tris, 150 mM NaCl, 1% TritonX-100, pH 7.6).

**Western blot**

4× Laemmli buffer (8% SDS, 40% glycerol, 20% β-mercaptoethanol, 0.01% bromophenol blue, and 250 mM Tris HCl pH 6.8), pH-adjusted with 1.5 M Tris HCl pH 8.8, was added to P2 synaptosomes or primary cultures at a 1x final concentration. Samples were boiled at 95 °C for 5 min, and loaded in a 4-12% polyacrylamide gels (Invitrogen). Protein was transferred into a nitrocellulose membrane using the semi-dry Trans-Blot® Turbo™ Transfer System (Biorad 1704150). Total protein was quantified using the REVERT Total Protein Stain Kit (Licor LI 926-11010). 5% milk solution in TBS-T (150 mM NaCl, 20 mM Tris, 0.5% Tween) was used to block and to prepare primary and secondary antibodies. SuperSignal West Femto Maximum Sensitivity Substrate (Thermo scientific 1859290) was used to develop westernblots. Image J was used to quantify intensities within regions of interest (ROI). For total protein quantification, intensity was averaged across 3 ROIs per lane, and ROI masks were placed at the same position for all lanes.

**ELISA detection of soluble and insoluble Aβ$_{42}$**

Hippocampi from $App^{NL}$ or $App^{NL\text{-}G\text{-}F}$ mice at 1, 2, 3 and 6 months were dissected after transcardial perfusion with ice-cold PBS. Tissue was homogenized in protein extraction reagent (Pierce). Homogenates were centrifuged at 4 °C for 1 h at 100,000×g (Beckman TLA 100.4 rotor) and supernatants used for ELISA. Guanidine-HCl extraction protocol was used to extract GuHCl-soluble Aβ fraction. Aβ42 levels were quantified on Meso Scale Discovery (MSD) 96-well plates by ELISA using end specific antibody provided by Dr. Marc Mercken (Janssen Pharmaceutica, Belgium). Monoclonal antibody JRFcAβ42/26 against the C terminus of Aβ42 species was used as capture antibodies and JRF/AβN/25 labeled with sulfo-TAG was used as the detection antibody. The plate was read in MSD Sector Imager 6000.

**Sleep deprivation**

At beginning of light phase/sleep cycle sleep deprivation (SD) starts. Animals are kept awake by gentle touch with a brush. At 3h and 4h of SD a novel object was inserted in each cage. Gentle poke with brush continues until 6h of SD. Group W: animals were perfused at the beginning of the light phase; Group SD: animals undergo SD from the beginning of light cycle for 6h and are perfused; Group SD + RB: animals undergo SD from the beginning of light cycle for 6h, are allowed to rebound sleep (RB) for 4h, and are perfused.

**Statistical analysis**

Data was plotted in GraphPad Prism 8. For quantification, datasets were tested for normality using D'Agostino and Pearson test. If datasets passed the test, they were analysed using Student's unpaired t test. Otherwise, the datasets were analysed using nonparametric unpaired t tests Mann–Whitney. One-way ANOVA test was used for multiple comparisons.

8. Oostrum, M. Van, Campbell, B., Müller, M. & Pedrioli, P. G. A. Surface proteome dynamics during neuronal development and synaptic plasticity. (2019).
